# Harnessing the Cross-talk between Tumor Cells and Tumor-associated Macrophages with a Nano-drug for modulation of Glioblastoma Immune Microenvironment

**DOI:** 10.1101/170282

**Authors:** Tong-Fei Li, Ke Li, Chao Wang, Xin Liu, Yu Wen, Yong-Hong Xu, Quan Zhang, Qiu-Ya Zhao, Ming Shao, Yan-Ze Li, Min Han, Naoki Komatsu, Li Zhao, Xiao Chen

## Abstract

Glioblastoma (GBM) is the most frequent and malignant brain tumor with a high mortality rate. The presence of a large population of macrophages (Mφ) in the tumor microenvironment is a prominent feature of GBM and these so-called tumor-associated Mφ (TAM) closely interact with the GBM cells to promote the survival, progression and therapy resistance of the GBM. Various therapeutic strategies have been devised either targeting the GBM cells or the TAM but few have addressed the cross-talks between the two cell populations. The present study was carried out to explore the possibility of exploiting the cross-talks between the GBM cells (GC) and TAM for modulation of the GBM microenvironment through using Nano-DOX, a drug composite based on nanodiamonds bearing doxorubicin. In the in vitro work on human cell models, Nano-DOX-loaded TAM were first shown to be viable and able to infiltrate three-dimensional GC spheroids and release cargo drug therein. GC were then demonstrated to encourage Nano-DOX-loaded TAM to unload Nano-DOX back into GC which consequently emitted damage-associated molecular patterns (DAMPs) that are powerful immunostimulatory agents as well as indicators of cell damage. Nano-DOX was next proven to be a more potent inducer of GC DAMPs emission than doxorubicin. As a result, Nano-DOX-damaged GC exhibited an enhanced ability to attract both TAM and Nano-DOX-loaded TAM. Most remarkably, Nano-DOX-damaged GC reprogrammed the TAM from a pro-GBM phenotype to an anti-GBM phenotype that suppressed GC growth. Finally, the in vivo relevance of the in vitro findings was tested in animal study. Mice bearing orthotopic human GBM xenografts were intravenously injected with Nano-DOX-loaded mouse TAM which were found releasing drug in the GBM xenografts 24 h after injection. GC damage was evidenced by the induction of DAMPs emission within the xenografts and a shift of TAM phenotype was detected as well. Taken together, our results demonstrate a novel way with therapeutic potential to harness the cross-talk between GBM cells and TAM for modulation of the tumor immune microenvironment.

**Abbreviations:** ATP, adenosine triphosphate; BBB, blood-brain barrier; BCA, bicinchoninic acid; BMDM, bone marrow derived macrophages; CD, cluster of differentiation; CFSE, 5(6)-carboxyfluorescein diacetate, succinimidyl ester; CM, conditioned culture medium; CNS, central nervous system; CRT, calreticulin; DAMPs, damage-associated molecular patterns; DAB, diaminobenzidine; DOX, doxorubicin; ECL, enhanced chemiluminescence; ELISA, enzyme-linked immunosorbent assay; HMGB1, high mobility group protein B1; HSP90, heat shock protein 90; FACS, flow cytometry; GBM, glioblastoma; Guanylate Binding Protein 5 (GBP5); GC, glioblastoma cells; IHC, immunohistochemical; IL, interleukin; Mφ, macrophages; mBMDM, mouse BMDM; mBMDM2, Type-2 mBMDM; M1, Type-1 Mø; M2, Type-2 Mø; Nano-DOX, ND-PG-RGD-DOX; ND, nanodiamonds; Nano-DOX-mBMDM, Nano-DOX-loaded mouse BMDM; NGCM, Nano-DOX-treated-GC-conditioned medium; PBS, phosphate buffered saline; PG, polyglycerol; PMA, phorbol 12-myristate 13-acetate; PVDF, polyvinylidene fluoride; RGD, tripeptide of L-arginine, glycine and L-aspartic acid; RM, regular culture medium; SD, standard deviation; TAM, tumor-associated Mφ; TBST, Tris Buffered Saline with Tween® 20.

**Graphic abstract:** 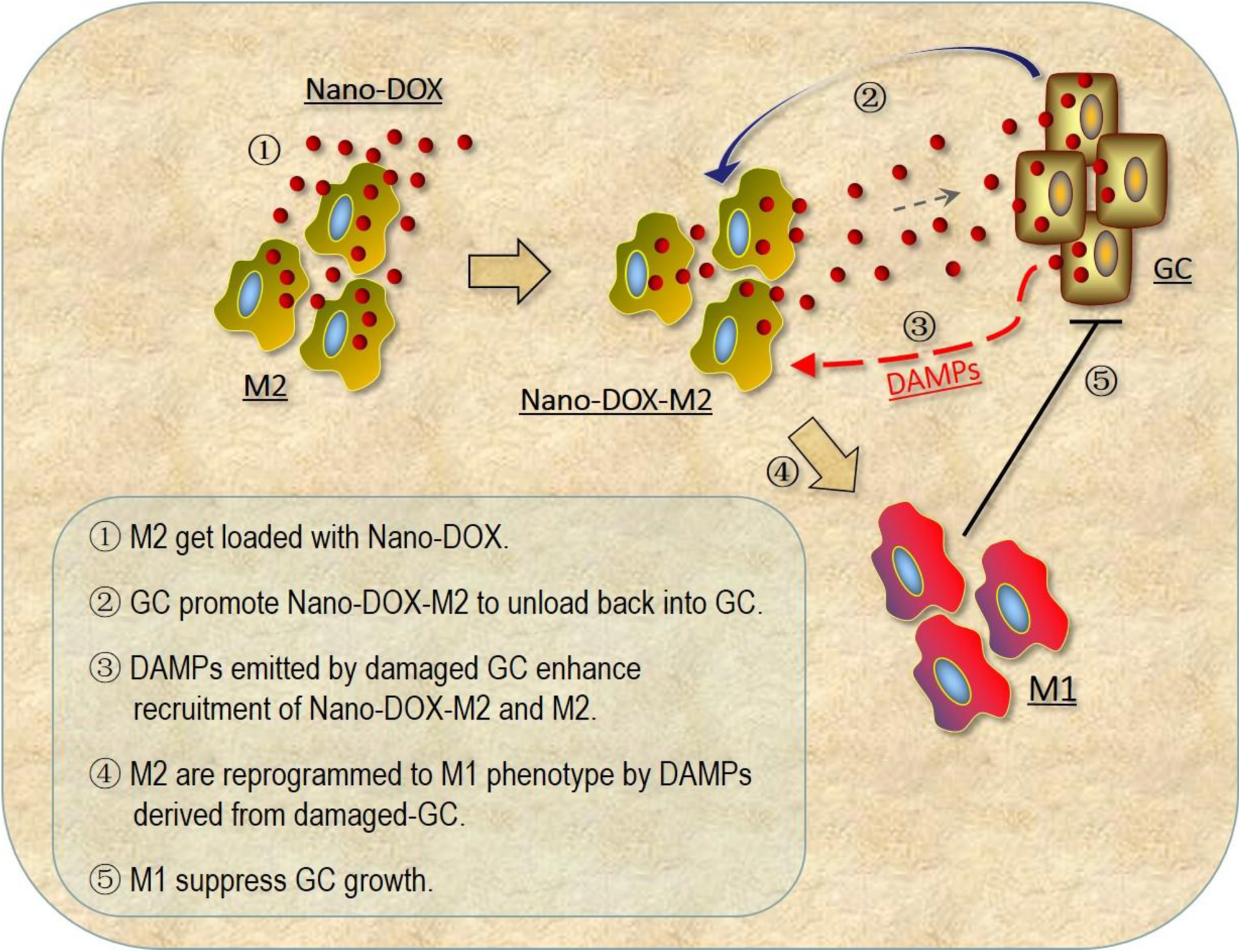

## 1. Introduction

Glioblastoma (GBM) is the most common and most deadly type of malignant primary brain tumor in adults [1]. Chemotherapy is indispensable in the treatment of all GBM patients but GBM has anatomical, pathological and metabolic features e.g. the blood-brain barrier (BBB), abnormal vasculature, a high interstitial hydrostatic pressure and an acidic extracellular pH that severely restrict the access of most chemotherapy drugs to the tumor tissue [2-4]. Various drug delivery strategies have been devised aiming to improve the drug availability in the GBM, the majority of which are designed for targeting the malignant cancer cells [5, 6]. However, as with other types of solid tumor, the GBM tissue is a heterogeneous ecosystem consisting not only of malignant cells but also a variety of non-malignant stromal cells like macrophages (Mø), fibroblasts, myofibroblasts, lymphocytes and endothelial cells, which have been revealed to closely interact with the malignant cells and provide active support for tumor survival, growth, progression and development of resistance to therapy [7, 8]. With increasing appreciation of their vital roles in the pathophysiology of cancers including GBM, the tumor stromal cells have received growing interests as therapeutic targets. But surprisingly, little is known about 1) how the drug delivery devices would affect the malignant cells’ dialogues with the stromal cells, and 2) how the stromal cell components in the GBM would interact with the drug delivery devices and affect their therapeutic responses.

In GBM, the most prominent members of tumor stromal cells are the so-called tumor-associated Mø (TAM) [9]. TAM in the GBM are largely recruited and derived from monocytes in the blood circulation and can reach as high as 40 percent of the tumor cell mass [10]. Monocytes recruited in diseased tissues differentiate into Mø that are activated into one of two broad phenotypes (i.e. type 1 or type 2) in response to local microenvironmental cues [11, 12]. Type-1 Mø (M1), also termed classically activated Mø, are immunostimulatory and characterized by induction of inducible nitric oxide synthase (iNOS), secretion of pro-inflammatory cytokines (e.g. IL-6, IL-12, IL-1β and TNF-α), upregulation of cell surface molecules for antigen presentation (e.g. MHC II, CD80 and CD86) and an enhanced phagocytic ability. M1 coordinate the creation of an adverse local inflammatory microenvironment for cancer cells and play central roles in the initiation and maintenance of anti-tumor immunity. Type-2 Mø (M2), also termed alternatively activated Mø, on the contrary, are anti-inflammatory and characterized by the production of immunosuppressive cytokines (e.g. IL-10, IL-1ra and TGF-β), downregulation of cell surface molecules for antigen presentation, upregulation of cell surface antigens FasL and B7-H1 which are necessary for induction of lymphocyte cell death. M2 suppress anti-tumor immunity and coordinate remodeling of the tumor microenvironment into one that favors tumor survival, growth, and progression. Ample evidence has suggested that TAM within the GBM are predominantly of the pro-tumor M2 phenotype [13-15]. Fundamental understanding of the importance of monocytes and TAM in GBM has encouraged great interest in manipulating them for therapeutic purposes. Proofs have emerged for the concept of using monocytes and Mø as active carriers overcoming the BBB to deliver chemotherapy drugs in GBM [16, 17]. Strategies to deplete TAM or modulate TAM phenotype and function have also shown therapeutic potential in the treatment of GBM [18, 19]. However, the roles as well as mechanisms of TAM-cancer cell interplay in achieving these promising outcomes still largely remain in the black box.

To shed light on the above-raised questions, the present study was carried out wherein the cross-talks between GC and TAM were mechanistically probed in response to Nano-DOX, a chemotherapy drug delivery device previously reported by us [20]. This device was based on functionalized nanodiamonds bearing doxorubicin (DOX), a first-line chemotherapy drug for treatment of multiple cancers. The composition of Nano-DOX is described in the “Material and methods” section and illustrated in Fig. 1. Nano-DOX is without any special mechanism for penetrating the BBB that protects GBM against access of foreign drug molecules. The use of Nano-DOX in the setting of GBM thus may seemingly lack practical relevance. Under inflammatory conditions like the brain cancer, however, the BBB is largely disrupted favoring drug traffic into the tumor [21], which has been taken advantage of by many drug delivery strategies. More importantly, the massive recruitment of peripheral monocytes by the GBM provides an alternative drug delivery route into the GBM circumventing the BBB [5, 22]. There have been pre-clinical study reports on monocyte-mediated drug delivery for treatment of glioma and depression [17, 23]. We in a separate study have also demonstrated that monocytes could deliver Nano-DOX in the GBM in a “Trojan horse” manner (submitted elsewhere). The delivered Nano-DOX are supposed to be taken up by the TAM as well as the GC. There has been study demonstrating uptake of therapeutic nanoparticles by TAM in a peripheral tumor model [24]. Thus, it is both relevant and coherent to use Nano-DOX in the current study. In vitro experiments were first performed on human GC and TAM models. The in vivo relevance of the in vitro findings was subsequently validated using mouse bone marrow-derived Mø (mBMDM) on mice bearing orthotopic human GBM xenografts.

**Fig. 1.**
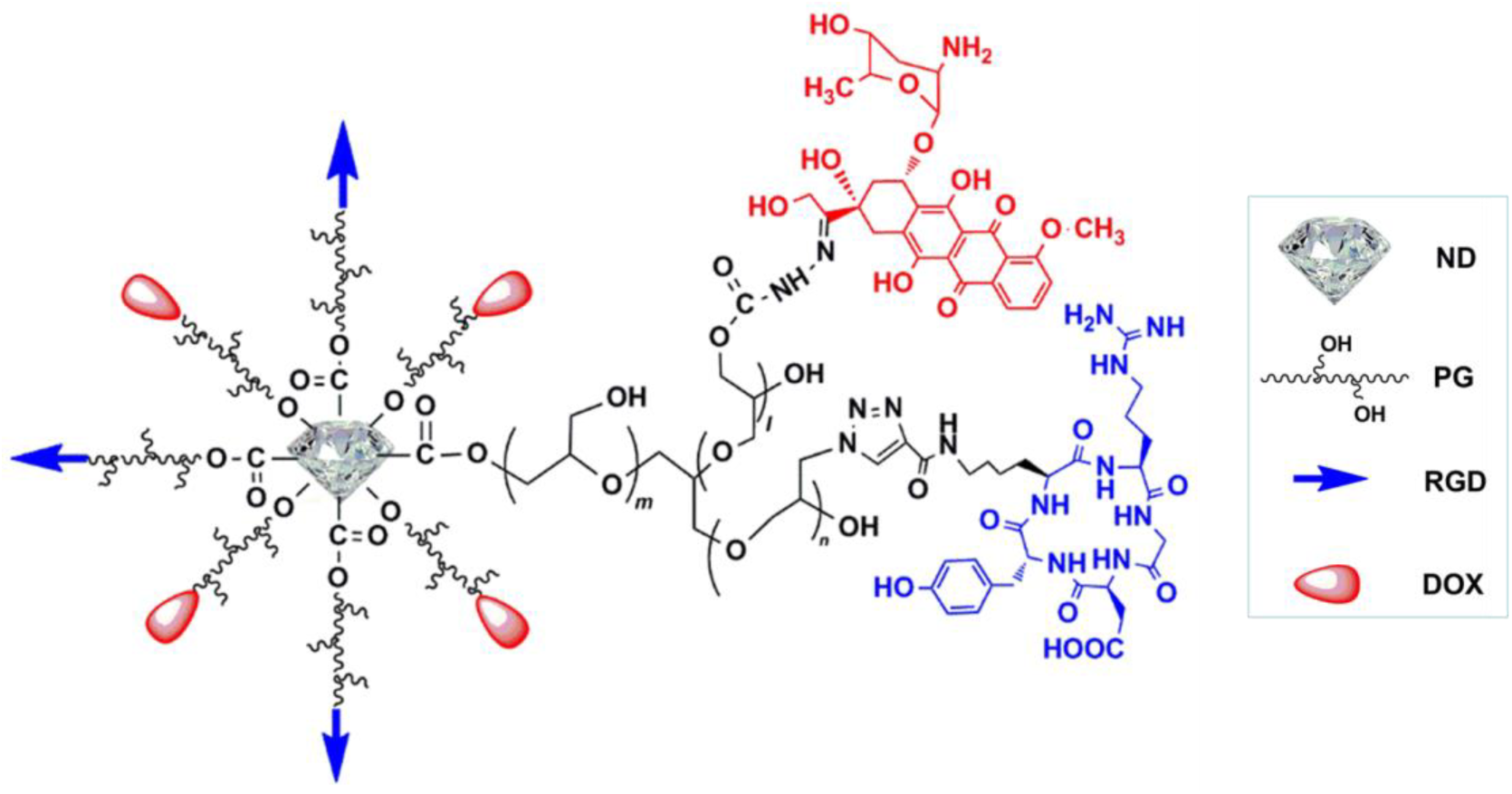
Composition and structure of Nano-DOX. ND: nanodiamond; PG: polyglycerol; RGD: cyclic tripeptide of L-arginine, glycine and L-aspartic acid; DOX: doxorubicin.

## 2. Material and methods

### 2.1 Cell Models

The Mø model in this study was differentiated from human histiocytic lymphoma U937 cells which are widely used in the biological study of MC and Mφ. Mouse bone marrow-derived Mø (mBMDM) were isolated according to a published protocol [25]. U87 MG cells, a human GBM cell line, were used as the brain tumor cell model and were referred to as GC in this work. Both cell lines were purchased from the Cell Bank of Shanghai Institutes for Biological Sciences (Shanghai, China). All cells were cultured in RPMI-1640 medium (Sigma-Aldrich, USA) supplemented with 10% fetal bovine serum (Sigma-Aldrich) in a humidified incubator (5% CO_2_/95% air atmosphere at 37℃) which was referred to as regular culture condition. U937 cells were differentiated into Mø (M0) through incubating with 100 ng/ml of phorbol 12-myristate 13-acetate (PMA) for 48h. M0 were next stabilized for 24 h in the fresh culture medium. Type-2 activation of Mø and (mBMDM) were achieved by incubating the cells with 20 ng/ml of interleukin-4 (IL-4, Peprotech) and 20 ng/ml of interleukin IL-13 (IL-13, Peprotech) for 48 h. Type-1 Mø activation was obtained by incubation of Mø with 20 ng/ml of interferon-γ (IFN-γ, Peprotech) and 100 ng/ml of lipopolysaccharides (LPS, Peprotech) for 48 h [26, 27]. Type-2 Mø and mBMDM were referred to as M2 and mBMDM2, respectively and type-1 Mø referred to as M1 hereafter. M2 and mBMDM2 phenotype was characterized by cell surface markers and cytokine secretion (Fig. S1 & S2).

### 2.2 Nano-DOX

This device was based on nanodiamonds (4-5 nm in diameter) with a surface coating of polyglycerol (ND-PG). ND-PG was attached with cyclic tripeptides of L-arginine, glycine and Laspartic acid (RGD) which serve as a targeting moiety owing to specific binding to the integrin receptor α_v_β_3_ that is over-expressed in multiple types of cancer cells including GC [28]. The NDPG-RGD composite was loaded with doxorubicin (DOX), a first-line chemotherapy drug for treatment of multiple cancers, to give the final product Nano-DOX. Nano-DOX has a hydrodynamic diameter of 83.9 ± 32.3 nm in water and has good solubility in physiological solutions. Nano-DOX was prepared and fully characterized in our previous work [20]. The composition of Nano-DOX is illustrated in Fig. 1. Nano-DOX stock solution in water was kept at 4 ℃ and was sonicated in a water bath for 3 min before being diluted with culture medium into working concentrations. All concentrations and dosages of Nano-DOX used in this study were normalized to DOX.

### 2.3 Drug uptake and cell viability assay

M2 in 96-well plates with a density of 8×10^3^ cells/well were treated with Nano-DOX and DOX at concentrations up to 4 μg/mL, 100 μL per well for 24 h. The cells were then photographed with a fluorescent microscope (IX73, Olympus, Japan) and cell viability was assayed with a CCK-8 kit as instructed in the manual provided by the kit manufacturer (Dojindo Molecular Technologies, Inc., Japan). On the basis of the cell viability assay results, the Nano-DOX-loaded M2 (Nano-DOX-M2) used in all experiments hereafter were prepared by incubating M2 with culture medium containing 2.5 μg/ml of Nano-DOX for 24 h unless noted otherwise.

### 2.4 Three-dimensional (3-D) GC spheroid infiltration assay

3-D GC spheroids were prepared according to a published protocol [29]. Briefly, a 96-well plate was pretreated with 80 μl of 2% (w/v) agarose gel to prevent cell adhesion. GC were seeded into the plate at a density of 2×10^3^ cells/well and cultured under regular condition for 7 days to allow spheroid formation. GC spheroids were then co-cultured with Nano-DOX-M2 or Nano-DOX-loaded mBMDM2 (Nano-DOX-mBMDM2) pre-labeled with 5(6)-carboxyfluorescein diacetate, succinimidyl ester (CFSE, Sigma-Aldrich, USA) at a density of 8×10^4^ cells/well. CFSE labeling was performed prior to Nano-DOX loading according to a published protocol [30]. After 12 h, the cell spheroids were rinsed with PBS, fixed with 4% paraformaldehyde, and transferred to a chambered coverslip for confocal microscopy.

### 2.5 Assay of intracellular drug content in mixed cell culture

Nano-DOX-M2 (2×10^5^ cells/well) were mixed-cultured with CFSE-labeled GC (2×10^5^ cells/well) in 24-well plates under regular culture condition for 24 h. Nano-DOX-M2 were also single-cultured as a control. Cells in mixed and single cultures were taken at 0 and 24 h and measured by flow cytometry (FACS) for intracellular drug content and CFSE staining as well. Nano-DOX-M2 and GC were also mix-cultured on coverslips using the above protocol as well for fluorescence visualization. Single-cultured Nano-DOX-M2 were also observed for comparison. Mixed and single cultures were visualized at 0 and 24 h with a confocal microscope (OLYMPUS, FV1200) after Hoechst 33342 nuclear counterstaining and 4% paraformaldehyde fixation.

### 2.6 Assay of DAMPs emission from GC under the influence of Nano-DOX-M2

GC without CFSE labeling were mixed-cultured with Nano-DOX-M2 on coverslips for 24 h as described above. Cell surface exposure of CRT was then detected by immunofluorescent staining. GC were also cultured in Nano-DOX-M2-conditioned medium for 24 h; culture medium supernatants were then collected and HMGB1 levels were determined by enzyme-linked immunosorbent assay (ELISA) and ATP levels determined by chemiluminescence assay. Nano-DOX-M2-conditioned medium was the supernatants of culture medium in which Nano-DOX-M2 (2×10^5^ cells/0.5 mL/well, 24-well plates) had been maintained for 24 h. M2-conditioned medium was used as a control. Levels of HMGB1 and ATP in Nano-DOX-M2-conditioned medium and M2-conditioned medium were assayed as described above and deducted from the final culture medium supernatants at 24 h.

### 2.7 Assay of DAMPs emission from GC treated with Nano-DOX or DOX

GC in 24-well plates with a seeding density of 2.5×10^5^ cells/well were treated with either Nano-DOX or DOX over a concentration range of 1 ~ 10 μg/mL for 24 h. Cell surface exposure of CRT and HSP90 was then detected by immunofluorescent staining, confocal microscopy, and FACS. Culture medium supernatants were collected and HMGB1 levels were determined by ELISA and ATP levels determined by chemiluminescence assay. Cellular HMGB1 expression was also analyzed by Western blotting analysis.

### 2.8 Transwell chemotaxis assay

Transwell devices (insert pore size 8.0 μm, Corning, 3422) were used for the experiments as illustrated in Fig. 7 A & B. GC were seeded in the lower chamber at a density of 1×10^5^ cells/well and M2 or Nano-DOX-M2 were seeded in the insert at a density of 5×10^4^ cells/well. After 18 h of co-culture, the insert was taken out, fixed with 4% paraformaldehyde and stained with crystal violet. After washing with phosphate buffer solution (PBS), the cells were observed using a fluorescent microscope and cells in each of 5~8 fields of view were counted.

### 2.9 Assay of drug content in Nano-DOX-M2 in mixed culture under the influence of Nano DOX-damaged GC

Nano-DOX-M2 (2×10^5^ cells/well) were either mixed-cultured with CFSE-labeled GC (2×10^5^ cells/well) or single-cultured in 24-well plates in Nano-DOX-damaged-GC-conditioned medium (NGCM) and, for comparison, regular culture medium (RM) and RM containing 1.72 μg/mL of Nano-DOX for 24 h. Cells in mixed and single cultures were taken at 0 and 24 h and measured by FACS for intracellular drug content. NGCM was prepared as follows. GC in 24-well plates were maintained in RM containing 2.5 μg/mL of Nano-DOX (2×10^5^ cells/0.5 mL/well) for 24 h. Supernatants of the culture medium were then collected as the NGCM which contained 1.72 ± 0.12 μg/mL of Nano-DOX.

### 2.10 Assay of phenotypic change in M2 under the influence of Nano-DOX-damaged GC

M2 in 24-well plates (2×10^5^ cells/well) were treated with NGCM, RM or RM containing 1.72 μg/mL of Nano-DOX for 24 h. The cells were then harvested for 1) Western blotting analysis of GBP5 expression, 2) analysis of cell surface markers CD80 and CD86 by immunofluorescent staining and FACS, and 3) extraction of total mRNA for real-time polymerase chain reaction (RT-PCR) analysis of gene expression of iNOS and IκB-α which reflects the transcription activity of NF-κB [31]. Culture medium supernatants were collected for ELISA analysis of cytokine levels of IL-6, IL-1β, and IL-10. In addition, the phagocytic function of the treated cells was assayed using carboxylate modified polystyrene yellow-green fluorescent latex beads (2.5%, diameter 1 μm, Sigma-Aldrich) and FACS according to a previously published protocol [20]. Fluorescent images were also acquired using a laser scanning confocal microscope.

### 2.11 Evaluation of GC proliferation under the influence of reprogrammed M2

In the first approach as illustrated in Fig. 10 A, co-culture was performed in a Transwell (insert pore size 0.4 μm, Corning, 3413) wherein GC were placed in the lower chamber with a seeding density of 1×10^5^ cells/well and M2 were seeded in the insert at a density of 8×10^4^ cells/well. After stabilization for 24 h, the co-culture was treated with Nano-DOX (2.5 μg/mL) for another 24 h before GC proliferation was assayed using a CCK-8 kit. For comparison, co-culture without Nano-DOX treatment and single culture of GC in the presence of Nano-DOX were also performed. In the second approach as illustrated in Fig. 10 C, CFSE-labeled GC were mixed-cultured with M2 in the presence of 1.72 μg/mL of Nano-DOX or NGCM containing 1.72 μg/mL of Nano-DOX for 24 h. The cells were taken out at 0 h and 24 for FACS analysis of CFSE staining, the decay of which is proportional to the rate of cell proliferation.

### 2.12 Immunofluorescent staining of surface-exposed CRT and HSP90 and Mø Surface markers

Cell cultured on coverslips were washed with PBS, stained with Hoechst33342 (5 μg/mL), fixed with 4% paraformaldehyde and blocked with 5% bovine serum albumin (BSA) solution at 37 ℃ for 1 h. The samples were then incubated with anti-human CRT antibody (ab92516, Abcam) or anti-human HSP90 antibody (ab13495, Abcam) at 4 ℃ overnight and further stained with Alexa Flour 488-conjugated secondary antibody (SA00006-2, Proteintech) at 37 ℃ for 90 min. For Mø Surface marker analysis, the samples were incubated with antibodies for CD80 (anti-CD80 FITC antibody, 11-0809, eBioscience), CD206 (anti-CD206 Alexa flour 488, 5s-2069, eBioscience), CD 86(anti-CD86 Alexa flour488, 53-0869, eBioscience) or CD163 (anti-CD163 APC, 326510, Biolegend) for 40 min at 4 ℃. Cells were imaged using a laser scanning confocal microscope. Alternatively, cells cultured in 24-well plates were harvested, subjected to the above staining protocol and analyzed by FACS.

### 2.13 Assay of released HMGB1, ATP and cytokines

HMGB1 levels in the cell culture medium supernatants were determined using a HMGB1 ELISA kit (Elabscience, E-EL-H1554c) and spectrometer (Biotek ELX800). Supernatant ATP levels were determined using a Chemiluminescence ATP Determination Kit (Beyotime, S0027, China) and a luminometer (Tecan, Spark 10M). IL-1β, IL-6 and IL-10 levels in culture medium supernatants were determined using ELISA kits (CHE0001, 4Abio; EK1062, EK1102, MultiSciences) and a spectrometer.

### 2.14 FACS analysis

Cellular fluorescence was acquired on a Cytoflex flow cytometer equipped with Beckman Coulter Cytoflex. CFSE, FITC and Alexa flour 488 fluorescence was acquired in the FL-1 channel. DOX and Nano-DOX fluorescence was acquired in the FL-2 channel. At least 1×10^4^ cells/per sample were acquired for single cultures and 2×10^4^ cells/per sample acquired for mixed cultures. Geometric means (GM) were used to quantify the fluorescent intensity.

### 2.15 RT-PCR

Total RNA was extracted using an RNA extraction kit (TRIpure Reagent, Aidlab). mRNA contained in 1 μg of total RNA was reverse transcribed into cDNA using a transcriptor cDNA synthesis kit (TRUEscript RT MasterMix, Aidlab). Amplification of cDNA was performed using an SYBRGreen qPCR Master Mix kit (Aidlab). Primers (5′ to 3′) were the following: forward (CATCACCATCTTCCAGGAGCGAGA) and reverse (TGCAGGAGGCATTGCTGATGATCT) for GADPH, forward (GTCACCTACCACACCCGAGATG) and reverse (CGCTGGCATTCCGCACAA) for iNOS, forward (GCTGAAGAAGGAGCGGCTACT) and reverse (TCGTACTCCTCGTCTTTCATGGA) for IκB. IκB mRNA level was assayed to reflect the transcription activity of NF-κB. RT-PCR was performed using a Bio-RAD CFX Connect Optics Module and data were analyzed using Bio-RAD CFXmanager. See Fig. S9 for the melting curves and amplification curves.

### 2.16 Western analysis of HMGB1 and GBP5 expression

GC or M2 subjected to required treatments in six-well plates were rinsed twice with ice-cold PBS and lysed in RIPA buffer with 1% protease inhibitor cocktail. Cell lysates were cleared by centrifugation and protein concentration was determined using a BCA kit. Equal amounts of proteins were fractionated by SDS-PAGE and transferred to a PVDF membrane. The membranes were blocked with 5% fat-free milk in TBST and incubated with antibodies against GBP5 (13220-1-AP, Proteintech), HMGB1 (ab79823, Abcam) and GADPH overnight at 4 ℃. Protein bands were imaged using a horseradish peroxidase-conjugated secondary antibody and ECL and the films were exposed using a Bio Imaging system (Syngene).

### 2.17 Mouse glioma model in vivo

Female athymic BALB/c nude mice at 4–5 weeks of age (18 ~ 20 g) were purchased from Shanghai Laboratory Animal Center at the Chinese Academy of Sciences (Shanghai, China). Animal handling and experimental procedures were conducted in line with protocols approved by the Animal Care Committee at the Wuhan University. Mice were housed in a temperature-controlled environment with fresh water and rodent diet available at all times. All inoculations and administrations were performed under Nembutal anesthesia. Orthotopic GBM xenografts were established by slowly injecting GC (4×10^5^ cells/8 μl in PBS) into the right corpus striatum of a mouse with the aid of a stereotaxic apparatus. At day 23 post GC inoculation, the animals were randomly divided into 5 groups (4 mice per group) which received injections via the lateral tail vein. Animals in group “Nano-DOX-mBMDM2” received Nano-DOX-mBMDM2 suspended in PBS. The Nano-DOX-mBMDM2 were pre-labeled with CFSE and prepared by loading mBMDM2 with 2.5 μg/mL of Nano-DOX for 24 h. For a mouse with a body weight of 20 g, 6 × 10^6^ Nano-DOX-mBMDM2 in 200 μl of PBS were injected. The dosage normalized to DOX was calculated to be 0.4 mg/kg (body weight). Animals in groups “DOX” and “Nano-DOX” received PBS solution of DOX or Nano-DOX of the same dosage. Animals in group “mBMDM2” received mBMDM2 suspended in PBS and animals in group “control” only received PBS. After 24 h, in vivo fluorescence images were acquired using an in vivo imaging system (XTREME BI, BRUKER, USA). Excitation wavelength was 480 nm and emission wavelength collected was 600 nm. The animal were then sacrificed and vital organs were harvest and imaged. Cryosections (5 μm) of brain tumor tissues were prepared for fluorescent microscopy and paraffin sections were prepared for immunohistochemical (IHC) staining analysis.

### 2.18 IHC analysis

Antibodies for IHC analysis included rabbit anti-human CRT antibody (ab92516, Abcam), rabbit anti-human HMGB1 antibody (ab79823, Abcam), rabbit anti-human HSP90 antibody (ab13495, Abcam), mouse anti-mouse CD86 antibody (ab213044, Abcam), rabbit anti-mouse mannose receptor antibody (ab64693, Abcam), rabbit anti-mouse iNOS antibody (ab15323, Abcam) and rabbit anti-mouse GBP5 antibody (132201-AP, Proteintech). Paraffin sections (5 μm) were dewaxed and rehydrated, antigen repaired with sodium citrate for 20min, then incubated in 3% hydrogen peroxide for 10min at room temperature. The paraffin sections were then blocked with 5% BSA for 30 min, stained with antibodies overnight at 4 ℃, washed with PBS and stained with secondary antibody (PV-9000, ZSGB-BIO) for 1 h at 37 ℃. DAB (ZLI-9018, ZSGB-BIO) was applied for coloration for 5 min at room temperature. Hematoxylin was used to stain the nucleus.

### 2.19 Statistical analysis

All data were analyzed with Sigmastat software. Statistical differences between groups were analyzed using One-way analysis of variance (ANOVA).

## 3. Results

### 3.1. GC encourage Nano-DOX-loaded M2 to release Nano-DOX back into GC

The present study was developed from the basic hypothesis that GC would cross-talk with the drug-loaded TAM and promote drug unloading therefrom. To test this hypothesis, we differentiated the human U937 monocytic cells into Mø which were further activated into M2 as the model of TAM. The M2 were incubated with Nano-DOX at concentrations up to 4 μg/ml (normalized to DOX) for up to 48 h. As shown in Fig. 2 A & B, Nano-DOX-loaded M2 (Nano-DOX-M2) varied little in cell viability from control cells at 24 h and significant loss of cell viability only occurred at concentrations above 2 μg/ml at 48 h. In sharp contrast, M2 started to lose cell viability at 1 μg/ml of DOX at 24 h. Furthermore, Nano-DOX-M2 (fluorescently labeled with carboxyfluorescein diacetate succinimidyl ester (CFSE)) were shown to be capable of infiltrating three-dimensional (3-D) GC spheroids indicating that Nano-DOX-M2 were functionally viable (Fig. 2 C). In view of M2’s good tolerability to Nano-DOX, the Nano-DOXM2 used in the 3-D GC spheroid assay and all experiments thereafter were prepared by incubating M2 with culture medium containing 2.5 μg/mL of Nano-DOX for 24 h unless noted otherwise.

**Fig. 2.**
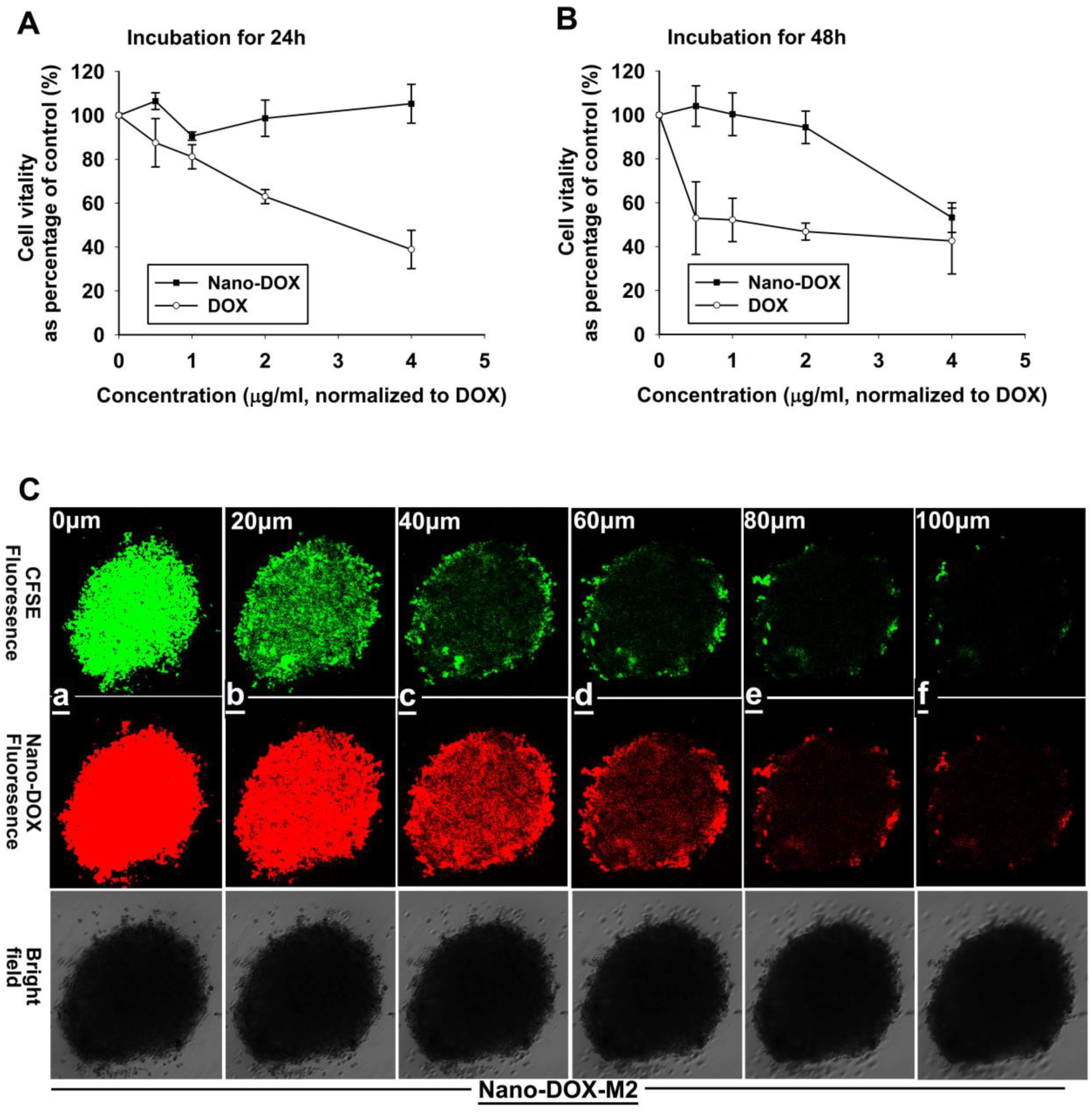
Functional viability of Nano-DOX-M2. A and B: M2 were treated with Nano-DOX or DOX for 24 h and 48 h and cell viability was assayed by CCK-8 test. Values were means ± standard deviation (SD) (n=4). C: Nano-DOX-M2 were mixed-cultured with GC spheroids for 12 h and the spheroids were photographed at different depths using a laser confocal microscope. Nano-DOX-M2 were prepared by incubating M2 with 2.5 μg/ml of Nano-DOX for 24 h.

To determine the influence of GC on drug unloading from Nano-DOX-M2, Nano-DOX-M2 were mixed-cultured with CFSE-labeled GC for 24 h before cellular Nano-DOX contents were measured by flow cytometry (FACS). As shown in Fig. 3 A-F, Nano-DOX-M2 both in the presence and absence of GC displayed reduced Nano-DOX fluorescence at 24 h indicating unloading of Nano-DOX-M2. However, Nano-DOX-M2 in mixed-culture with GC lost much more Nano-DOX fluorescence than in single culture indicating that GC promoted drug unloading from Nano-DOX-M2 (Fig. 3 F). In mixed culture, the loss of Nano-DOX fluorescence in Nano-DOX-M2 was accompanied by a gain of Nano-DOX fluorescence in GC indicating that GC took up the drug unloaded by Nano-DOX-M2 (Fig. 3 D-G). Interestingly but not surprisingly, Nano-DOX-M2 in mixed culture with GC also displayed a gain of CFSE fluorescence (Fig. 3 C, D & H) indicating that Nano-DOX-M2 received something from GC at the same time of giving away their cargo. It was also noted that decay of CFSE fluorescence in GC which is proportional to the rate of cell division was slower in the mixed culture than in single culture indicating suppressed GC proliferation by Nano-DOX-M2 (Fig. 3 B, D, E & I). Confocal microscopy provided visual evidence for Nano-DOX-M2 unloading into GC in mixed culture wherein GC featured an extended, spindle-like morphology that are readily distinguishable from the round-shape of Nano-DOX-M2 (Fig. 3 K). At 0 h, red fluorescence indicative of Nano-DOX was only seen in Nano-DOX-M2 but occurred in all cells after 24 h of mixed culture (Fig. 3 J & K). Concerning the nature of the drug unloaded by Nano-DOX-MC, Fig. 3 K and Fig. 4 A show that the drug unloaded from Nano-DOX-M2 entered primarily in the cytoplasm of GC. In single culture, free Nano-DOX stayed in the cytoplasm of GC as well whereas free DOX essentially went into the nuclei of GC (Fig. 5 A and Fig. 6 C). It is thus deduced that Nano-DOX-MC unloaded Nano-DOX but not DOX into GC. Taken together, the above observations strongly suggest that GC would encourage drug-loaded TAM to release their payload back into GC.

**Fig. 3.**
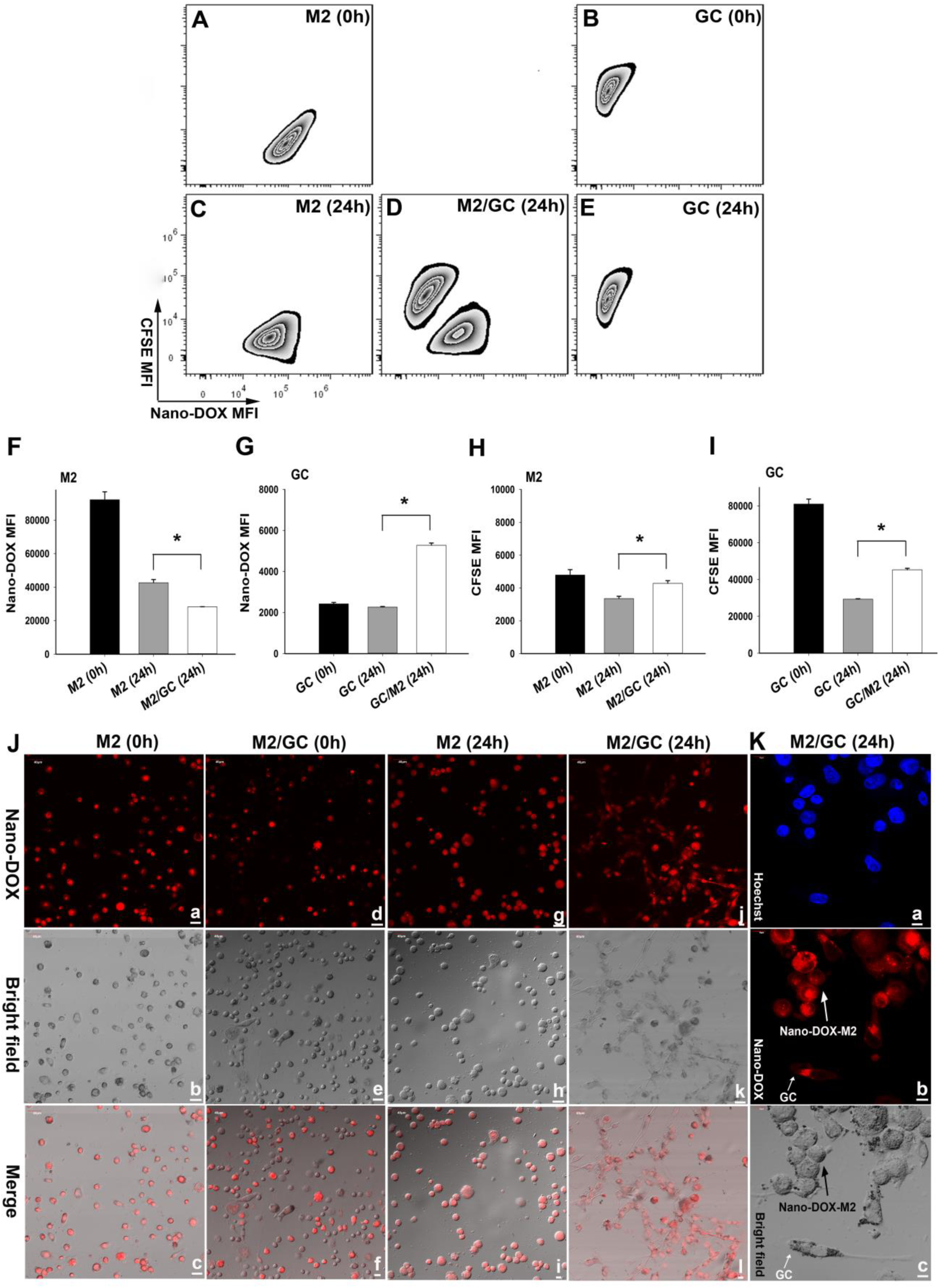
GC promote unloading of Nano-DOX-M2 back to GC. A-E: Nano-DOX-M2 were mixed-cultured with CFSE-labelled GC for 24 h. Single-cultured Nano-DOX-M2 served as control. Cellular fluorescence of Nano-DOX and CFSE was measured by FACS. F-H: Geometric means were used to quantify fluorescence intensity. Values were means ± SD (n=3, * p < 0.05). J: Cellular fluorescence of single-cultured Nano-DOX-M2 upon seeding (0 h, a-c); cellular fluorescence of Nano-DOX-M2 in single-culture for 24 h (g-i); cellular fluorescence of Nano-DOX-M2 in mixed culture with GC upon seeding (0 h, d-f); cellular fluorescence Nano-DOXM2 and GC in mixed-culture for 24 h (j-l). K: Subcellular distribution of fluorescence in Nano-DOX-M2 and GC in mixed-culture for 24 h. The observation was made with a confocal microscope. Red fluorescence comes from Nano-DOX and blue fluorescence is nuclear counterstaining.

**Fig. 4:**
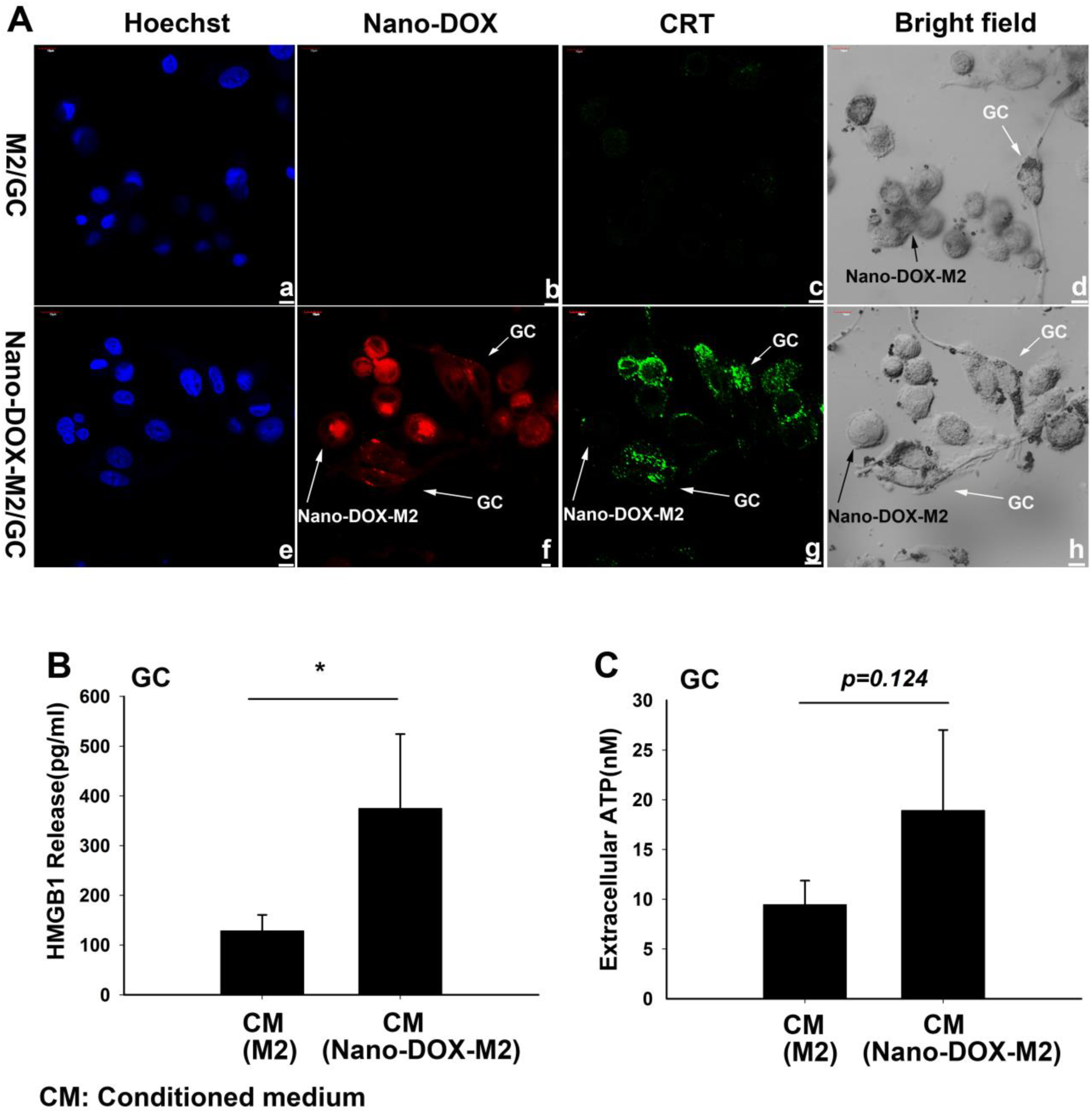
Nano-DOX-M2 induce DAMPs emission in GC. A: Nano-DOX-M2 or M2 were mixed-cultured with GC for 24 h. Cell surface exposure of CRT was detected by immunofluorescent staining. Green fluorescence is CRT staining; red fluorescence comes from Nano-DOX and blue fluorescence is nuclear counterstaining. B and C: GC were incubated with culture medium conditioned by Nano-DOX-M2 or M2 for 24 h and supernatant levels of HMGB1 and ATP were measured by ELISA and Chemiluminescence, respectively. Basal HMGB1 and ATP levels in the conditioned medium were subtracted. Values were means ± SD (n=3, * p < 0.05).

**Fig. 5.**
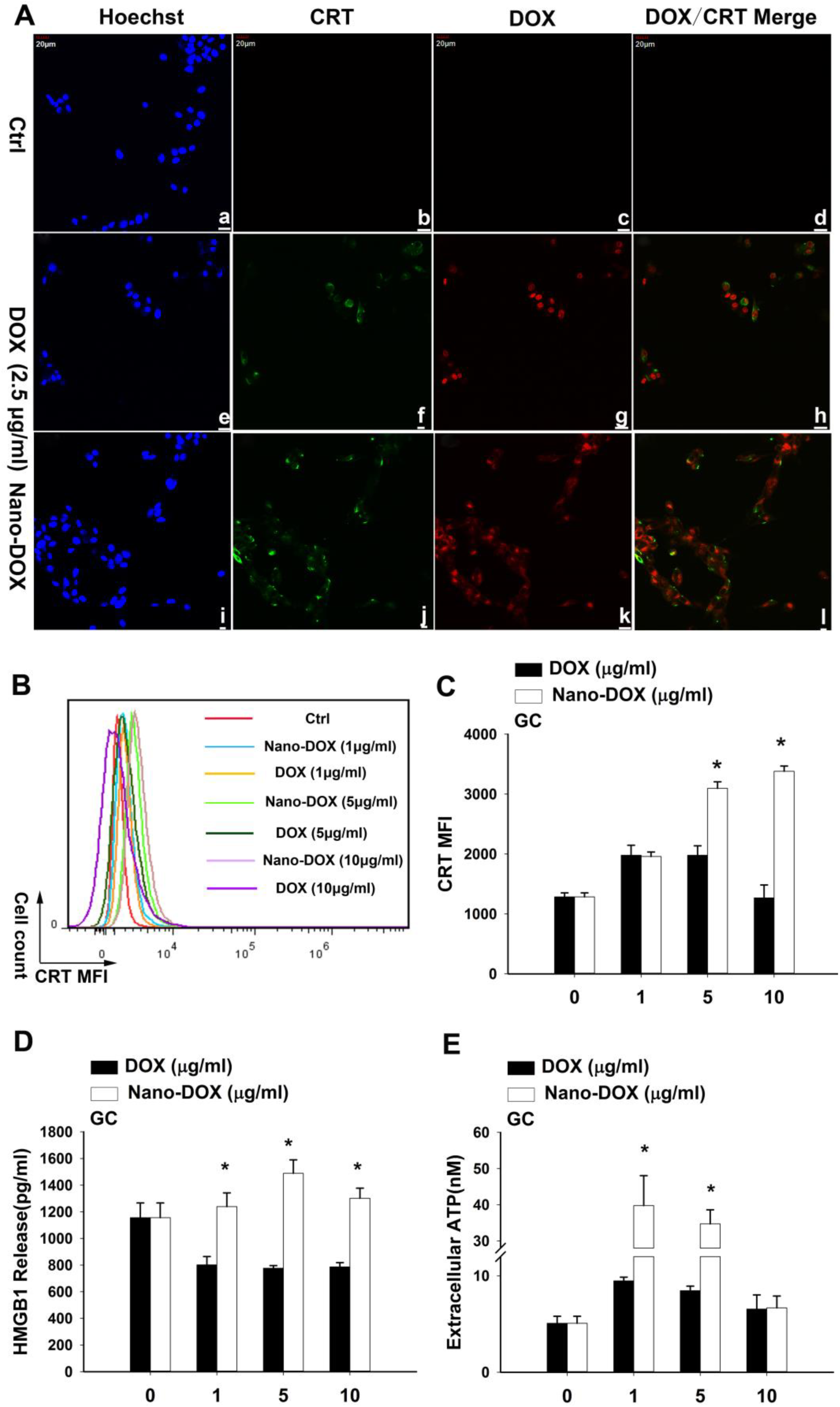
Nano-DOX is more potent than DOX in inducing emission of CRT, HMGB1 and ATP. GC were treated with Nano-DOX or DOX for 24 h before observation and analysis of DAMPs emission. A: confocal microscopy. Green fluorescence is CRT staining; red fluorescence comes from Nano-DOX or DOX and blue fluorescence is nuclear counterstaining. Images of triple merge of Hoechst, CRT and DOX fluorescence is provided in Fig. S5-C. B and C: FACS analysis of CRT emission. Geometric means were used to quantify fluorescence intensity. D and E: Supernatant levels of HMGB1 and ATP were measured by ELISA and Chemiluminescence, respectively. Values were means ± SD (n=3, * p < 0.05).

**Fig. 6.**
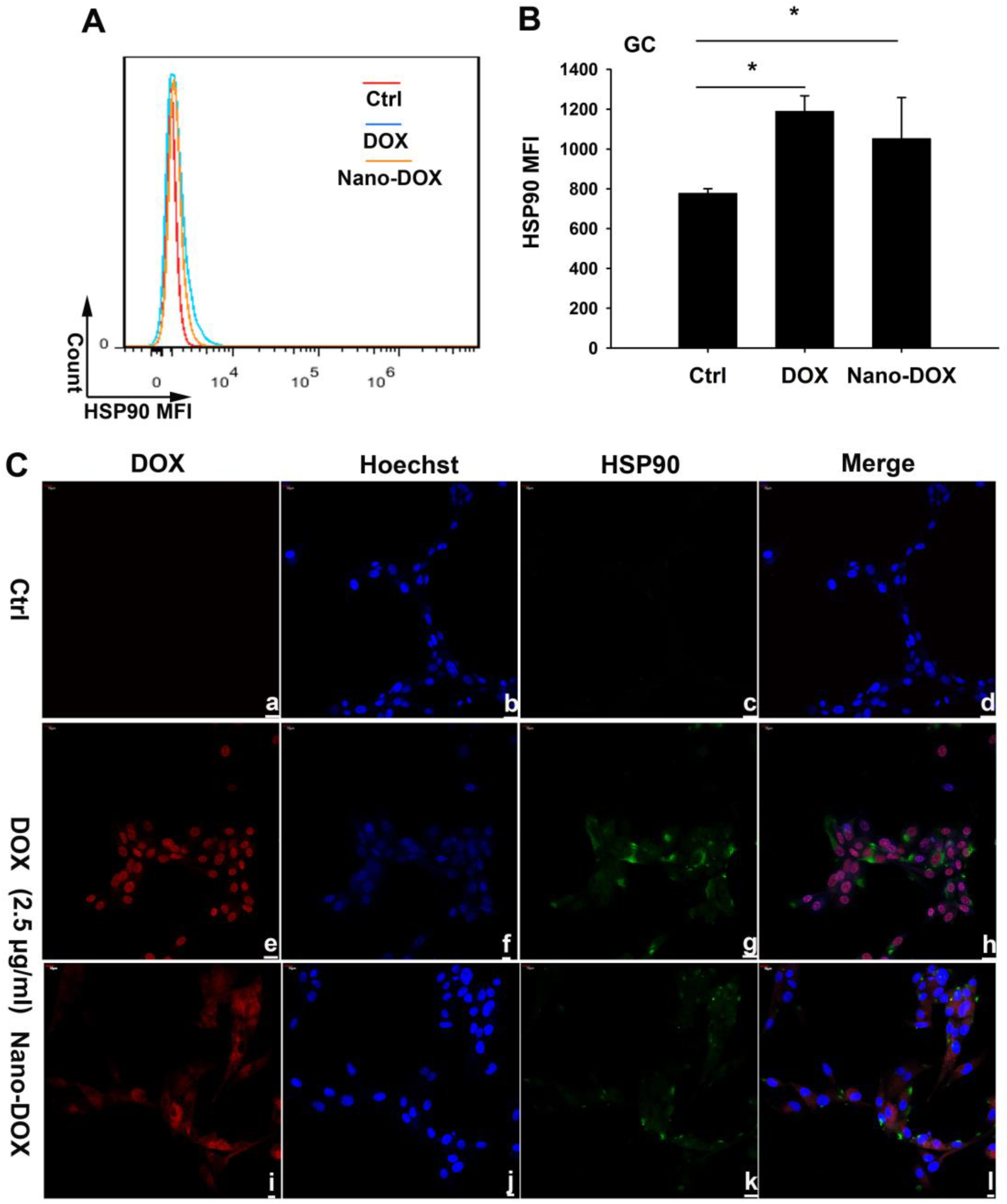
Nano-DOX is as potent as DOX in inducing emission of HSP90. GC were treated with Nano-DOX or DOX (2.5 μg/mL) for 24 h before observation and analysis of HSP90 emission. A and B: FACS analysis. Geometric means were used to quantify fluorescence intensity. Values were means ± SD (n=3, * p < 0.05). C: confocal microscopy. Green fluorescence is HSP90 staining; red fluorescence comes from Nano-DOX or DOX and blue fluorescence is nuclear counterstaining.

### 3.2. GC emit damage-associated molecular patterns (DAMPs) after taking up the Nano-DOX released by Nano-DOX-M2

In our previous study, the cytotoxicity of Nano-DOX manifested as suppression of cell proliferation [20]. This effect was also suggested in a later part of the present study (Fig. 10). Of greater importance, another line of evidence was found revealing GC damage inflicted by Nano-DOX released from Nano-DOX-M2. Cells in response to damage, stress or undergoing cell death are known to emit a spectrum of endogenous molecules termed damage-associated molecular patterns (DAMPs) [32-34]. Well established DAMPs include cell surface-exposed calreticulin (CRT), passively released high-mobility group box 1 protein (HMGB1) and actively secreted adenosine triphosphate (ATP). In our work, pronounced cell surface exposure of CRT in concurrence with Nano-DOX fluorescence were observed in GC that were mixed-cultured with Nano-DOX-M2 for 24 h (Fig. 4 A). A spike in HMGB1 release and a tendency toward higher secretion of ATP were also detected in GC that were incubated with Nano-DOX-M2-conditioned medium (Fig. 4 B, C). It is thus strongly indicated that Nano-DOX-M2 unloaded Nano-DOX into GC and induced GC damage. It is noted that Nano-DOX-M2, which were of a round shape distinct from the elongated, spindle-like morphology of GC, also displayed cell surface expression of CRT, both in mixed culture (Fig. 4 A) and single culture (Fig. S4)

### 3.3. Nano-DOX is a more potent inducer of DAMPs emission than DOX

Nano-DOX was originally designed to be able to shed DOX in the intracellular acid lysosomal compartment. As DOX is a well-demonstrated DAMPs inducer [35], it was not clear whether Nano-DOX or the DOX shed from Nano-DOX was responsible for the induction of DAMPs emission observed in GC (Fig. 4). To answer this question, we compared Nano-DOX and DOX (over a concentration range of 1-10 μg/ml) for their DAMPs-inducing activity in single-cultured GC. Nano-DOX overall displayed a higher potency than DOX in inducing emission of CRT, HMGB1, and ATP (Fig. 5) except in the case of heat shock protein (HSP90), another member of DAMPs, where Nano-DOX displayed no significant difference than DOX (Fig. 6). Surprisingly, DOX was found to reduce HMGB1 emission in GC (Fig. 5 D), which was corroborated by Western blotting analysis that showed decreased protein expression of HMGB1 in DOX-treated GC (Fig. S5 A) and the discrepancy between DOX and Nano-DOX was not due to any difference in their effects on cell proliferation (Fig. S5 B).

DAMPs perform predominantly non-immunological functions within the cell under physiological conditions. Under conditions of stress, damage or injury, these molecules are emitted to the cell surface or in the extracellular space becoming immunostimulatory danger signals [32-35]. In normal tissues, emitted DAMPs activate and perpetuate innate immunity for damage control. In solid tumor tissues, cancer cells are known to recruit various immune cells e.g. Mø into the tumor stroma to form a local immune microenvironment that provides aid and protection for tumor survival and progression [13-15]. When cancer cells are under attack for example by chemotherapeutic drugs, it would be natural of the tumor-associated stromal immune components to be mobilized by the DAMPs derived from the stressed, damaged or dying cancer cells. How the tumor-associated stromal immune components respond to cancer cell-derived DAMPs and their consequent reactions to cancer cells are heatedly explored questions of great importance both in cancer research and therapy. In this work, now that Nano-DOX-M2 have been demonstrated to incite DAMPs emission in GC, we next investigated the response of M2 to Nano-DOX-damaged GC and their consequent action back on GC.

### 3.4. Nano-DOX-damaged GC reinforce recruitment of M2 and Nano-DOX-M2

It is a logical assumption that more immune cells would be summoned to the tumor’s aid when tumor cells are damaged by Nano-DOX. One argument therefor is that certain DAMPs like ATP are “find-me” signals competent of promoting phagocyte migration [36]. The assumption was tested on a transwell co-culture model where either M2 or Nano-DOX-M2 were cultured in the upper insert with a bottom of membrane filter allowing cell passing through while GC were simultaneously cultured in the lower chamber as illustrated in Fig. 7 A & B. Indeed, both M2 and Nano-DOX-M2 in the inserts were attracted by GC to the underside of the bottom membrane in significantly higher numbers in the presence of Nano-DOX (2.5 μg/ml) than in absence (Fig. 7 C-F). Enhanced recruitment of TAM and Nano-DOX-TAM by Nano-DOX-damaged GC is thus suggested. We had initially also assumed that damaged GC could be more potent in stimulating unloading of Nano-DOX-M2. However, as shown in Fig. 7 H and Fig. S7, unloading of Nano-DOX-M2 was of the same extent in Nano-DOX-damaged-GC-conditioned medium (NGCM) as in the culture medium containing the same amount of Nano-DOX (1.72 μg/ml). But notably, GC were still found to stimulate unloading of Nano-DOX-M2 even in the presence of free Nano-DOX (Fig. 7 H, Fig. S7).

**Fig. 7.**
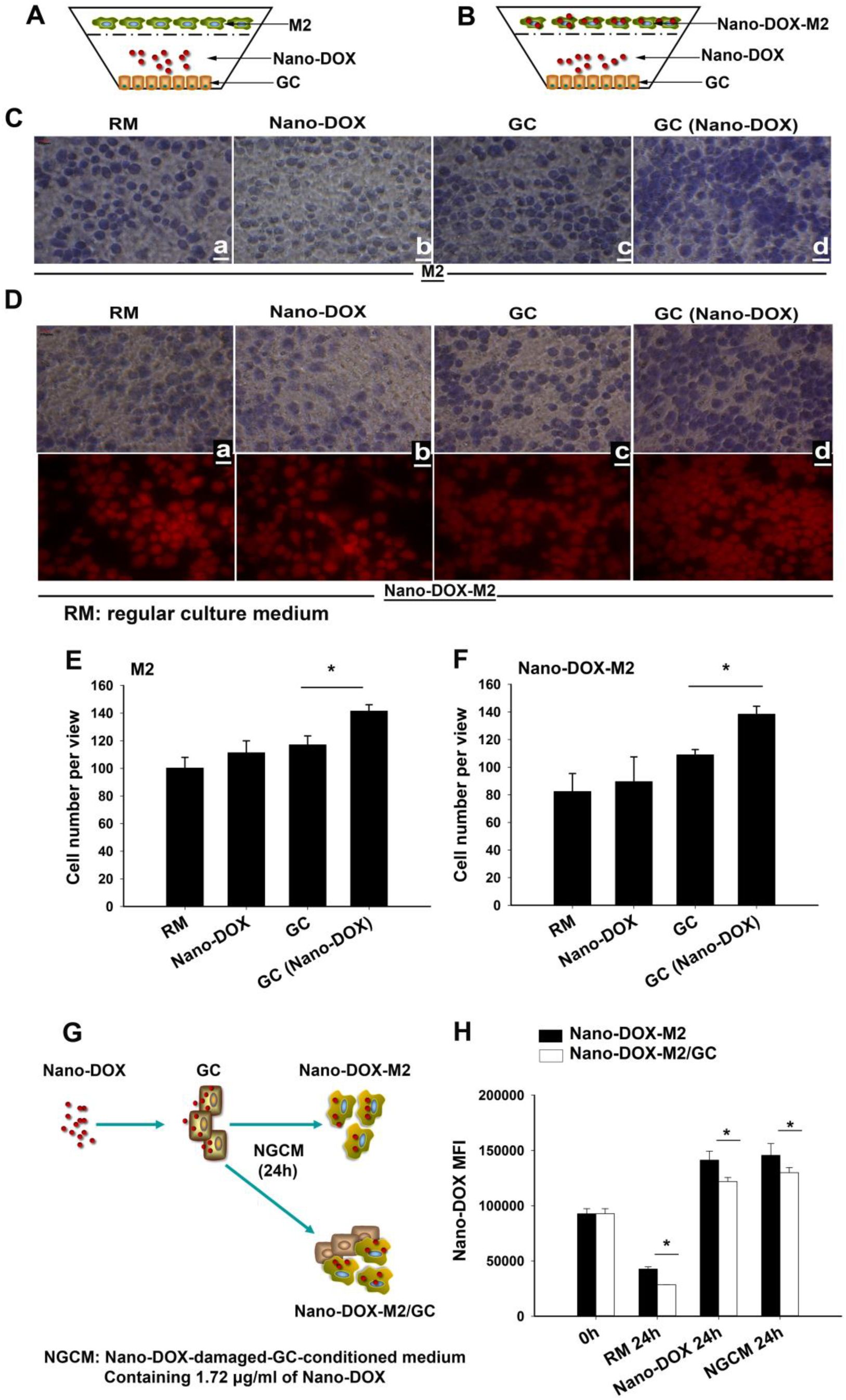
Nano-DOX-damaged GC reinforce recruitment of M2 and Nano-DOX-M2 but do not further promote unloading of Nano-DOX-M2. A and B: Nano-DOX-M2 or M2 were co-cultured with GC in the presence of 2.5 μg/mL of Nano-DOX in a Transwell for 18 h as illustrated. C and D: M2 or Nano-DOX-M2 that were attracted by GC to the underside of the insert bottom membrane were photographed. E and F: Cells in 5~8 fields of view were counted. G: Nano-DOX were mixed cultured with GC in the presence of NGCM for 24 h as illustrated and cellular fluorescence was measured with FACS at 0 h and 24 h. H: Geometric means were used to quantify fluorescence intensity. See Fig. S7 for representative FACS zebra dot plots. NGCM represents Nano-DOX-damaged-GC-conditioned medium containing 1.72 μg/mL of Nano-DOX. Values were means ± SD (n=3, * p < 0.05).

### 3.5. Nano-DOX-damaged GC reprogram M2 toward an anti-tumor M1-like phenotype

Since DAMPs are immunostimulatory danger signals competent of initiating innate immunity which involves primarily pro-inflammatory Mø activity [37-39], we supposed that DAMPs derived from Nano-DOX-damaged GC should be able to activate M2 into an M1-like phenotype and function. To test this supposition, M2 were incubated with NGCM (containing 1.72 μg/ml of Nano-DOX) for 24 h before being examined for phenotypic and functional changes (Fig. 8 A). As shown in Fig. 8 B-G, NGCM-treated M2 displayed remarkably elevated abundance of cell surface CD80 and CD86 and expression of GBP5, which are markers for type-1 macrophage activation. A significant increase in the secretion of IL-6 and IL-1β (cytokines characteristic of type-1 Mø activation) along with a tendency toward lower secretion of IL-10 (a cytokine characteristic of type-2 Mø activation) were also detected in NGCM-treated M2 (Fig. 9 A-C). Furthermore, NGCM-treated M2 displayed a heightened activity of NF-κB (quantified by IκB-α mRNA level), a key transcription factor whose activation is closely involved in type-1 Mø activation (Fig. 9 D & S8). Most remarkably, NGCM-treated M2 significantly upregulated gene expression of iNOS (Fig. 9 E & S8) and exhibited a markedly enhanced phagocytic activity (Fig. 9 F-H & S9-2) both of which are salient features of type-1 Mø activation. It was also noted that Nano-DOX itself appeared to have little effect on M2 phenotype. Finally, M2 under the influence of Nano-DOX-damaged GC were found to suppress GC proliferation, which was demonstrated in two approaches. In the first approach as illustrated in Fig. 10 A, GC growth was assayed by CCK-8 test after being co-cultured with M2 in a Transwell in the presence or absence of 1.72 μg/ml of Nano-DOX for 24 h. As shown in Fig. 10 B and Fig. S10, M2 promoted GC growth in the absence of Nano-DOX and Nano-DOX alone had little effect on GC growth. But when GC were together with M2 in the presence of Nano-DOX, GC growth was significantly suppressed. In the second approach as illustrated in Fig. 10 C, GC proliferation was assayed by the decay of cellular CFSE staining after being mixed-cultured with M2 in NGCM or culture medium containing 1.72 μg/ml of Nano-DOX for 24 h. As shown in Fig. 10 D & E, suppression of GC growth was appreciable when the mixed culture was treated with Nano-DOX but was more pronounced in the presence of NGCM. Together, these findings are compelling evidence that Nano-DOX-damaged GC reprogramed M2 toward an anti-GC M1-like phenotype.

**Fig. 8.**
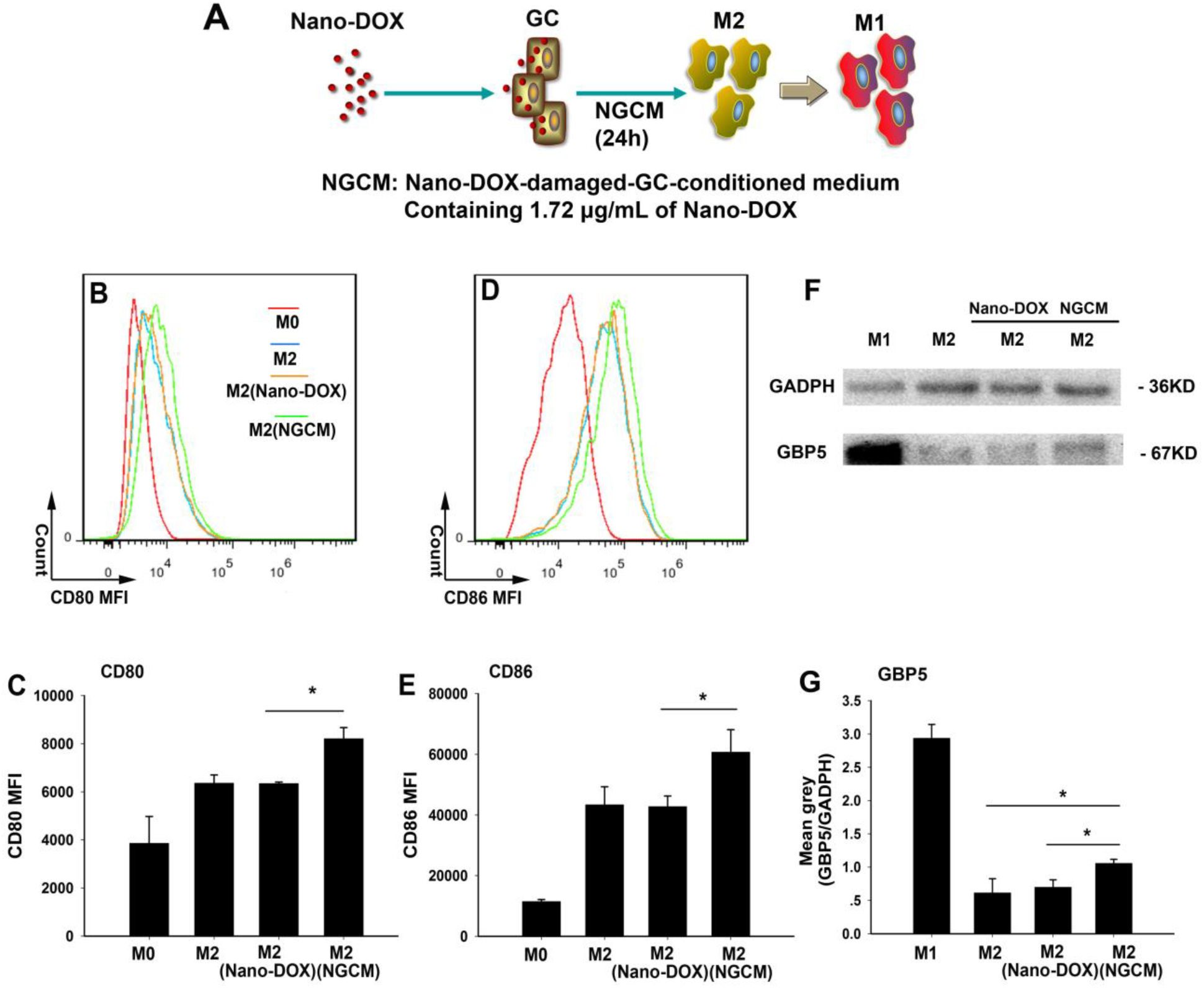
Nano-DOX-damaged GC reprogram M2 towards an M1-like phenotype. A: M2 were treated with NGCM for 24 h as illustrated. B-E: Cell surface markers CD80 and CD86 were analyzed by immunofluorescent staining and FACS. Geometric means were used to quantify fluorescence intensity. Values were means ± SD (n=3, * p < 0.05). Single-cultured undifferentiated Mø (M0) and Nano-DOX-treated M2 were used as controls. Increase of CD80 and CD86 expression in type-1activated macrophages (M1) relative to unactivated macrophages (M0) are presented in Fig. S9. F & G: GBP5 protein expression was assayed by Western blotting and blot bands was densitometrically analyzed using the software ImageJ (National Institutes of Health, USA).

**Fig. 9.**
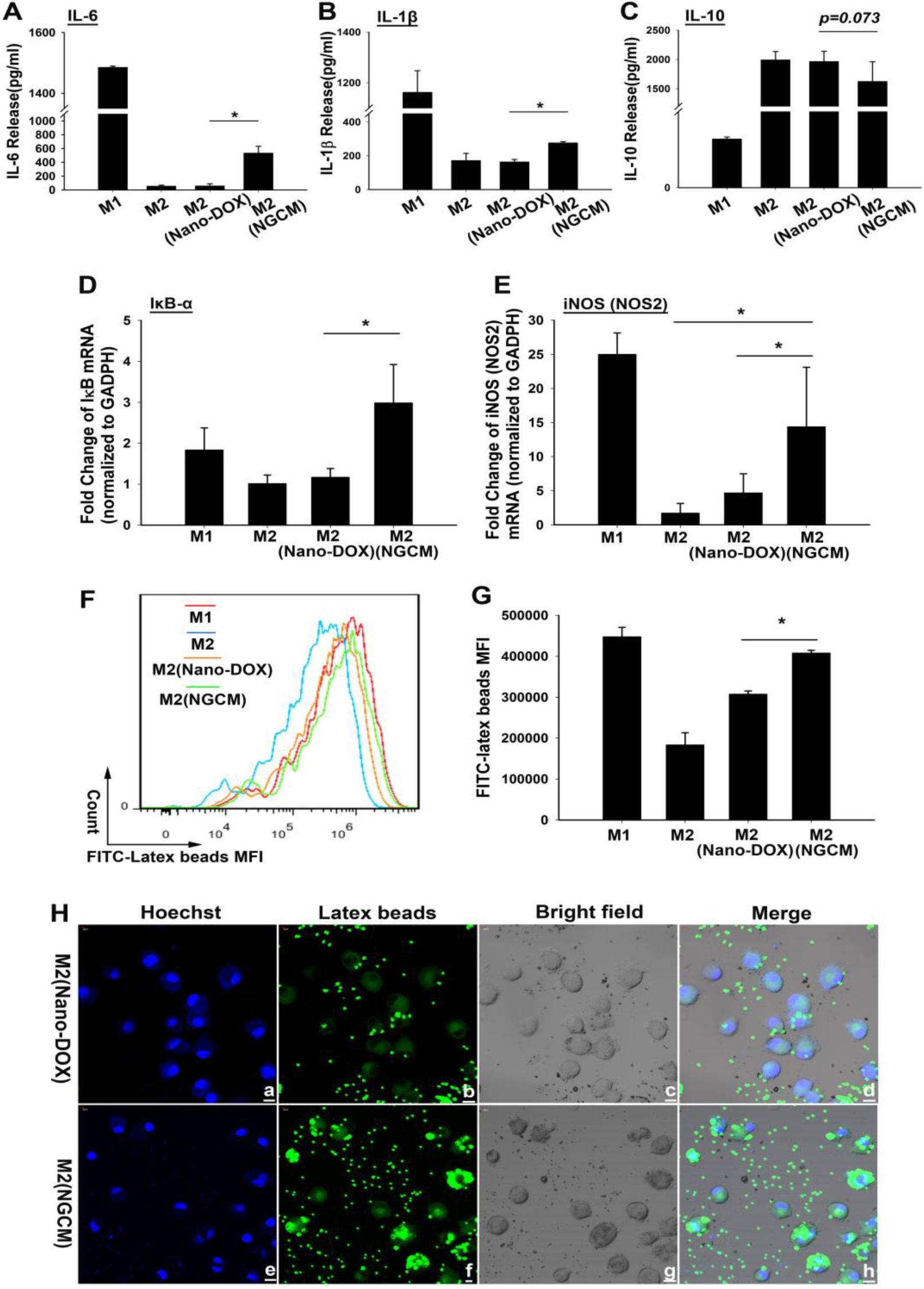
Nano-DOX-damaged GC reprogram M2 towards an M1-like phenotype. M2 were treated with NGCM for 24 h. A-C: Cytokine levels of IL-6, IL-1β and IL-10 were assayed by ELISA. D and E: The cells were harvest for analysis of mRNA levels of IκB-α and iNOS. F-H: Phagocytic ability was analyzed using fluorescent latex beads, FACS and confocal microscopy. Geometric means were used to quantify fluorescence intensity. Values were means ± SD (n=3, * p < 0.05). See Fig. S8 for the melting curves and amplification curves for D and E. See Fig. S9-2 for a more comprehensive version of H with more controls.

**Fig. 10.**
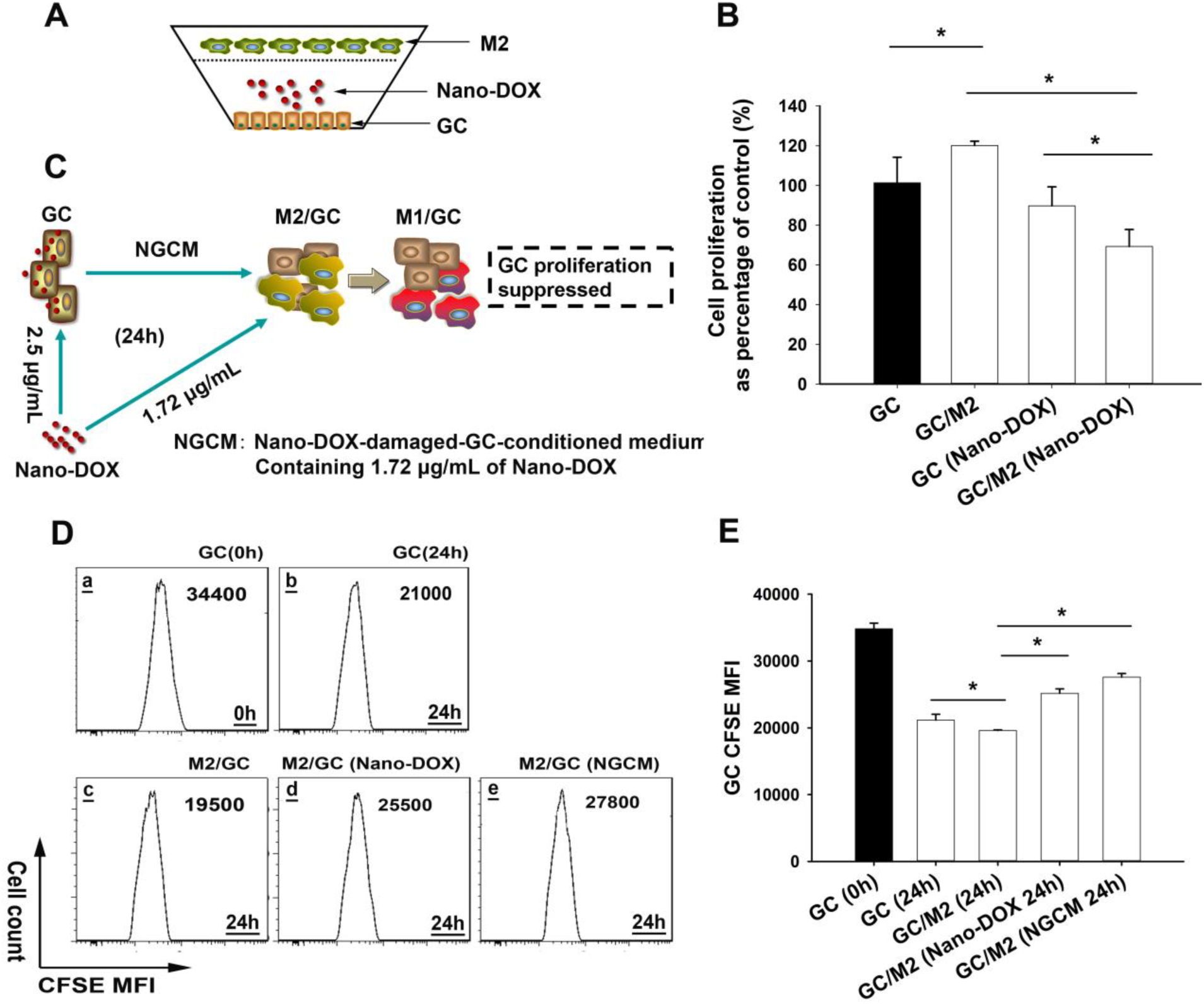
M2 reprogrammed by Nano-DOX-damaged GC inhibit GC proliferation, which was demonstrated in two approaches. A and B: Approach one wherein M2 were co-cultured with GC in the presence of 2.5 μg/mL of Nano-DOX in a Transwell for 24 h and GC proliferation was then evaluated by CCK-8 test. See Fig. S10 for visuals of cell growth upon the CCK-8 test. C-E: Approach two wherein CFSE-labeled GC were mixed-cultured with M2 in the presence of Nano-DOX or NGCM for 24 h. Cellular CFSE fluorescence was measured at 0 h and 24 h by FACS and decay of CFSE reflected the rate of GC proliferation. Values were means ± SD (n=3, * p < 0.05).

### 3.6. Nano-DOX-mBMDM2 release Nano-DOX in orthotopic GC xenografts inducing GC damage and Mø phenotype change

To validate the in vivo relevance of the in vitro findings, mBMDM were prepared and artificially polarized into the M2 phenotype. The type-2 mBMDM were then labeled with CFSE and loaded with Nano-DOX to yield Nano-DOX-mBMDM2. Functional viability of Nano-DOX-mBMDM2 was demonstrated by Transwell chemotaxis assay and 3-D GC spheroid infiltration assay (Fig. S11). Nano-DOX-mBMDM2 were intravenously injected into mice bearing orthotopic human GC xenografts. This experimental approach is based on the rationale that leukocytes have an intrinsic propensity to home to inflammatory sites e.g. the tumor tissue, which has been exploited for drug delivery in the brain [16, 17, 23]. In vivo and ex vivo imaging showed very weak DOX-derived fluorescence in the brains of xenograft-bearing mice 24 h after injection of DOX (Fig. 11 B & G) and Nano-DOX (Fig. 11 C & H) suggesting difficulty in accessing the brain for DOX and Nano-DOX. Mice injected with mBMDM2 displayed little fluorescence in the brains (Fig. 11 D & I). In sharp contrast, intensified fluorescence was observed in the brains of mice that had received Nano-DOX-mBMDM2 indicating delivery of Nano-DOX in the brain (Fig. 11 E & J). Certain vital organs e.g. the heart, liver and kidneys of mice injected with Nano-DOX or Nano-DOX-mBMDM2 also displayed various degrees of fluorescence suggesting possible drug accumulation at these sites (Fig. 11 H & J).

**Fig. 11.**
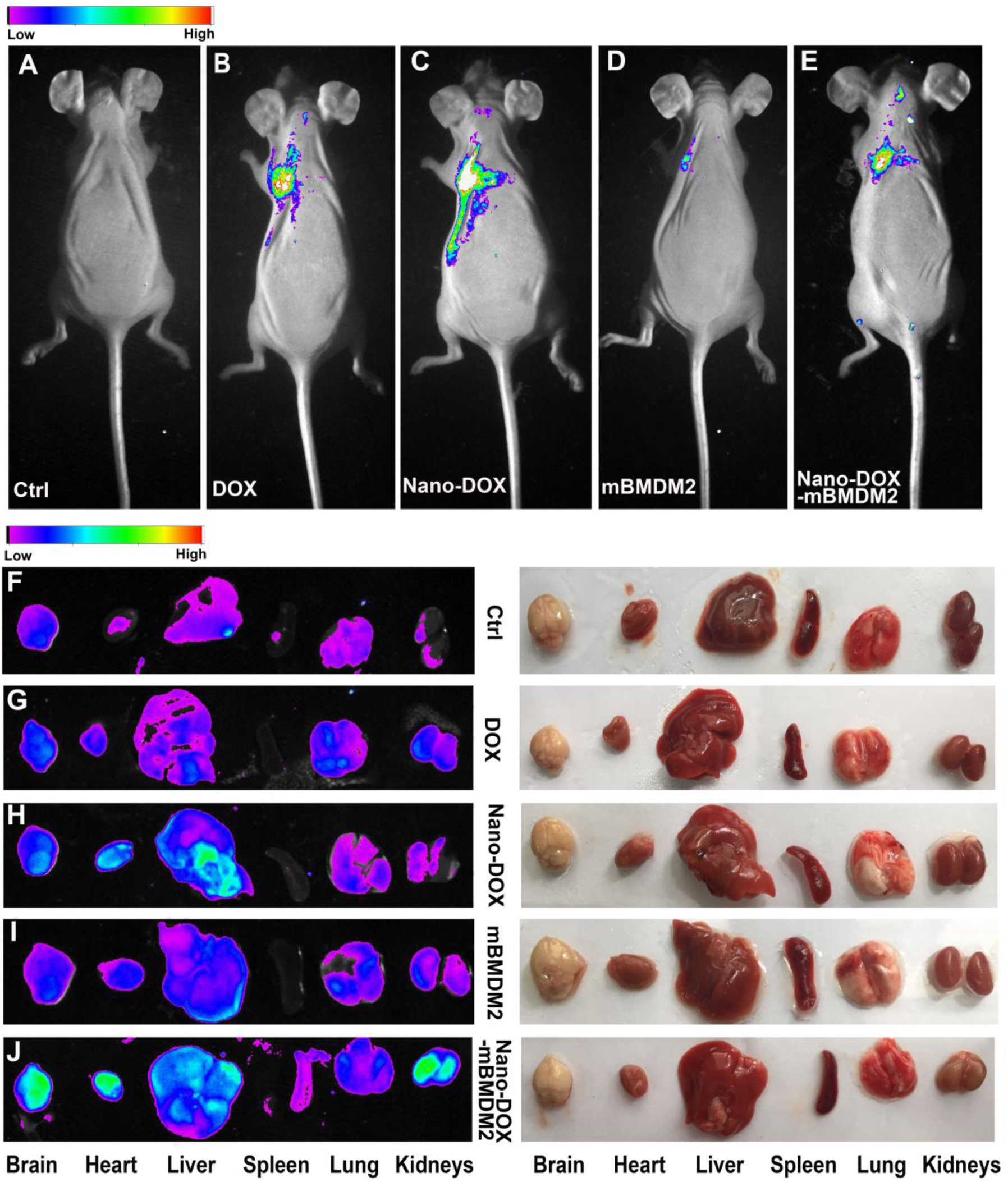
Drug distribution in mice bearing GC xenografts 24 h after intravenous injection of DOX, Nano-DOX, mBMDM2 and Nano-DOX-mBMDM2. A-E: In vivo imaging of drug fluorescence. F-J: Ex vivo imaging showing drug fluorescence in tumor xenografts and vital organs (left panel) and corresponding bright field imaging (right panel).

In brain tissue sections, GC were identified by tumor tissue structure, cell morphology and density (Fig. 12 and Fig. S12). In mice treated with Nano-DOX-mBMDM, green fluorescence from CFSE indicating the appearance of mBMDM2 and red fluorescence signifying Nano-DOX were both distinctly seen in the tumor xenograft tissue. (Fig. 12 M-O) The good agreement between the distribution patterns of green and red fluorescence strongly indicates the presence of Nano-DOX-mBMDM2 in the tumor tissue. The distribution of Nano-DOX fluorescence (red) was notably more extensive than the cell label fluorescence (green) indicating drug unloading from Nano-DOX-mBMDM2. In contrast, mice treated with DOX (Fig. 12 D-F) or Nano-DOX (Fig. 12 G-I) exhibited much weaker red fluorescence and no green fluorescence in the xenograft tissues suggesting much lesser brain-penetrating capability of the drugs per se. Mice injected with mBMDM2 (Fig. 12 J-L) only displayed the green fluorescence of CFSE evidencing the presence of mBMDM2 in the xenograft tissues.

**Fig. 12.**
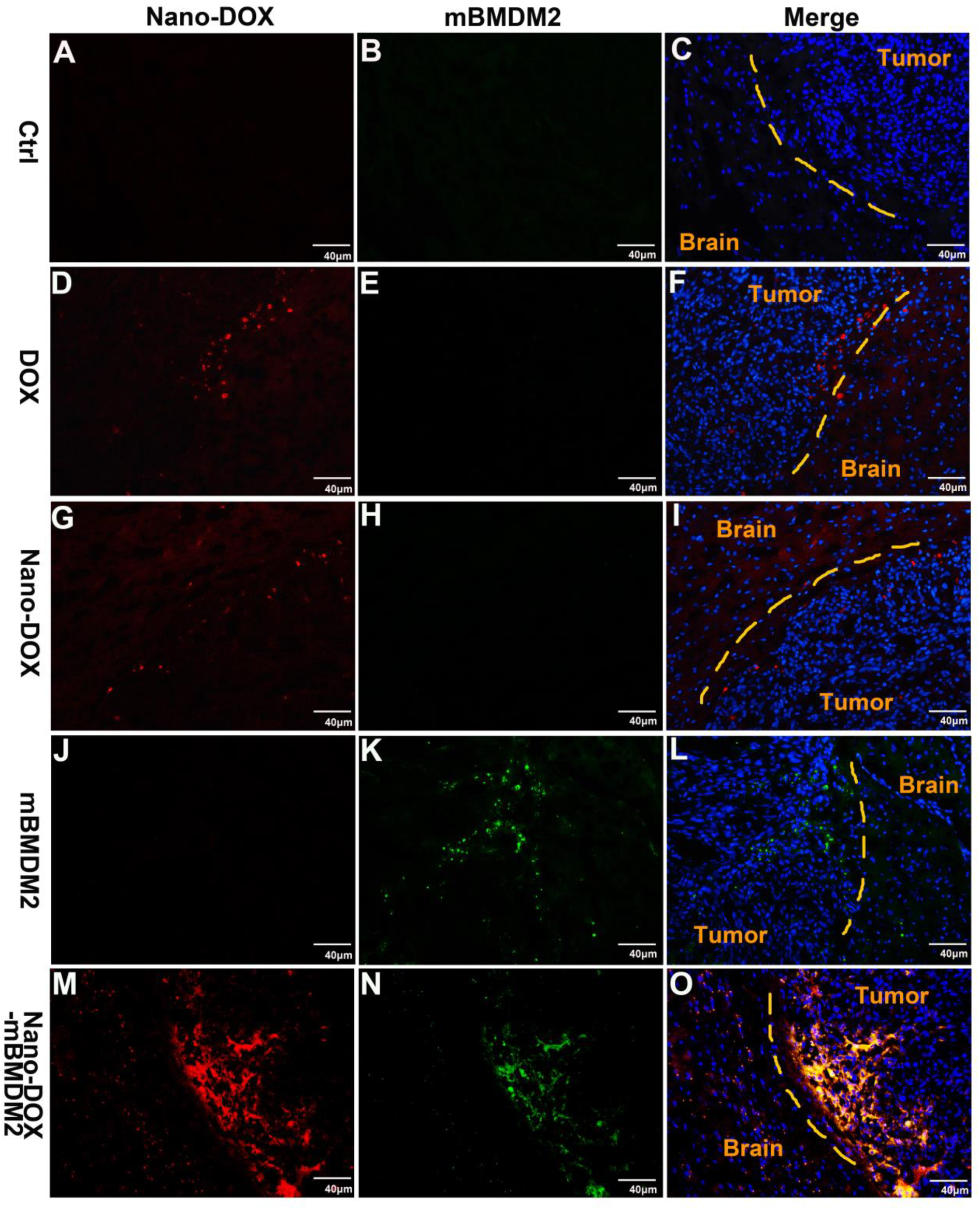
Fluorescent imaging of mice brain tissue sections shows the presence of Nano-DOX-mBMDM2 in orthotopic GC xenografts and release Nano-DOX therein. Blue fluorescence is nuclear staining of Hoechst 33342. Red fluorescence comes from DOX or Nano-DOX and green fluorescence from the CFSE that labels mBMDM2.

In consistence with the fluorescent observations, Nano-DOX-mBMDM2 induced markedly increased GC emission of DAMPs (CRT, HMGB1 and HSP90) over the other treatments in the xenograft tissues evidencing tumor cell damage (Fig. 13 E, J & O). By contrast, xenograft tissue DAMPs emission induced by DOX (Fig. 13 B, G & L) or Nano-DOX (Fig. 13 C, H & M) appeared to be comparable to control or slightly higher indicating little GC damage. Injection of mBMDM2 appeared to not increase DAMPs emission in the xenograft tissues (Fig. 13 D, I & N). Finally, immunohistological staining analysis showed that, in comparison with control (Fig. 14 A, F, K & P) and mBMDM2-injected mice (Fig. 14 D, I, N & S), mice that had received Nano-DOX-mBMDM2 (Fig. 14 E, J, O & T) exhibited increased expression of iNOS, GBP5, CD86 (markers of type-1 Mø activation) along with decreased expression () of CD206 (cell surface marker of type-2 Mø) in the brain tumor xenografts indicating the occurrence of a M2-M1 phenotype shift of Mø. DOX (Fig. 14 B, G, L & Q) or Nano-DOX (Fig. 14 C, H, M & R) overall did not elicit significant changes in the expression of these Mø markers except that higher-than-control GBP5 expression was observed with Nano-DOX treatment.

**Fig. 13.**
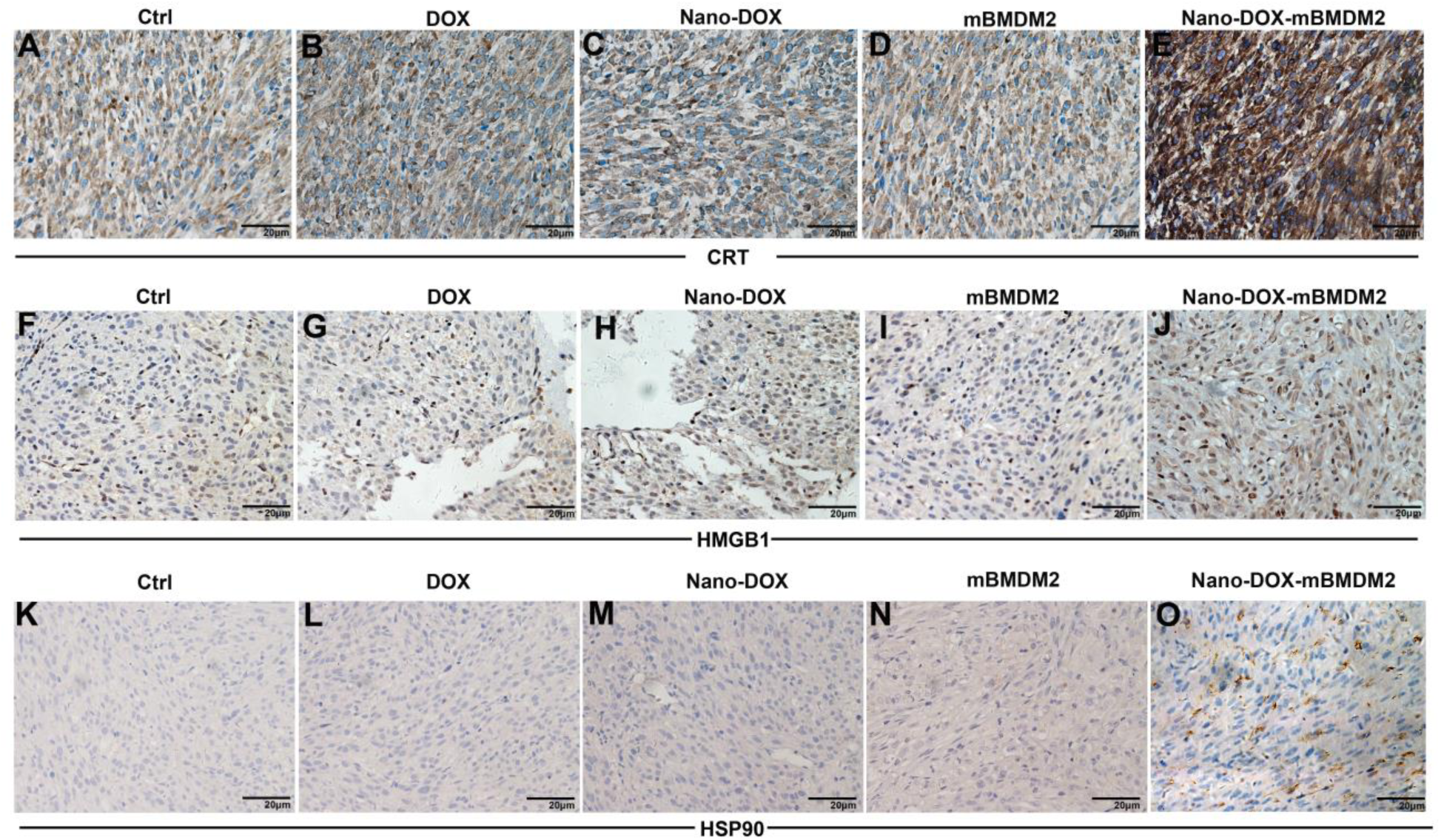
Immunohistological analysis shows markedly intensified or concentrated staining of DAMPs (CRT, HMGB1 and HSP90) in orthotopic GC xenograft tissues from mice treated with Nano-DOX-mBMDM2.

**Fig. 14.**
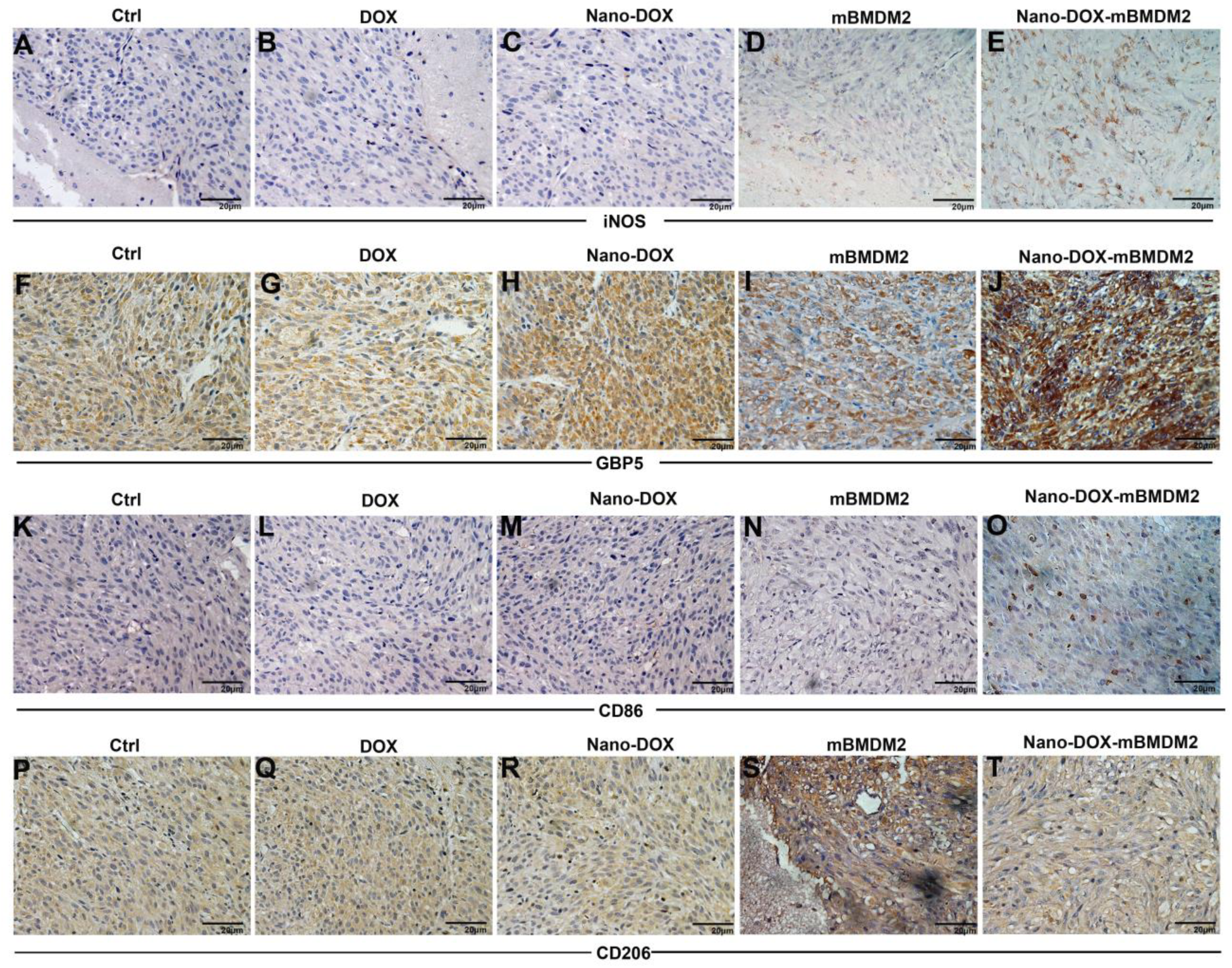
Immunohistological staining shows increased expression of iNOS, GBP5 and CD86 (markers of type-1 Mø activation) along with decreased staining of CD206 (marker of type-2 Mø activation) in orthotopic GC xenograft tissues from mice treated with Nano-DOX-mBMDM2.

Altogether, the above data strongly suggest that intravenously administered Nano-DOX-mBMDM2 could home to the GBM xenografts and release cargo drug therein and thereby damage the GC which in turn reprogram the mBMDM2 into a M1 phenotype.

## 4. Discussion

The interplay between malignant tumor cells and tumor stromal cells has been confirmed and investigated for its mechanisms from various angles by a lot of studies and has tremendous implication for tumor therapy. In the present work, we have drawn upon knowledge from these studies and demonstrated that GC have close cross-talk with their TAM, which could be exploited on two dimensions for therapeutic modulation of the GBM microenvironment (Fig. 15). First, peripherally recruited TAM which make up a high portion of the tumor mass could serve both as a drug carrier and reservoir to deliver chemotherapy drugs to the GC and cause cell damage. In this respect, Nano-based drug delivery devices (nano-drugs) may be advantageous over regular small-molecule drugs, most of which are difficult to penetrate the BBB and therapeutic concentrations can only be achieved at the cost of severe peripheral adverse effects [40, 41]. We clearly demonstrate the poor access of DOX to the brain and while Nano-DOX per se also has limited access to the brain, Mø could serve as an effective carrier to deliver Nano-DOX into the brain tumor (Fig. 11, Fig. 12). We also successfully used monocytes to deliver Nano-DOX into mouse orthotopic GC xenografts in another work (submitted elsewhere). On the contrary, DOX could not be carried by Mø or monocytes due to their high sensitivity to DOX’s toxicity (Fig. 2). Through mechanisms yet to be elucidated, Nano-DOX-loaded TAM (type-2 Mø) remain functionally viable and are stimulated to shed their drug cargo by the cancer cells (Fig. 2, Fig. 3 & Fig. 7). Preliminary findings suggest that DOX induces apoptosis whereas Nano-DOX is a potent inducer of autophagy which is probably implicated in the mechanisms by which Mø tolerate Nano-DOX (Data not shown). Secretory lysosomes and autophagosomes may be involved in the mechanisms whereby cancer cells promote unloading of Nano-DOX-loaded TAM (Data not shown).

**Fig. 15.**
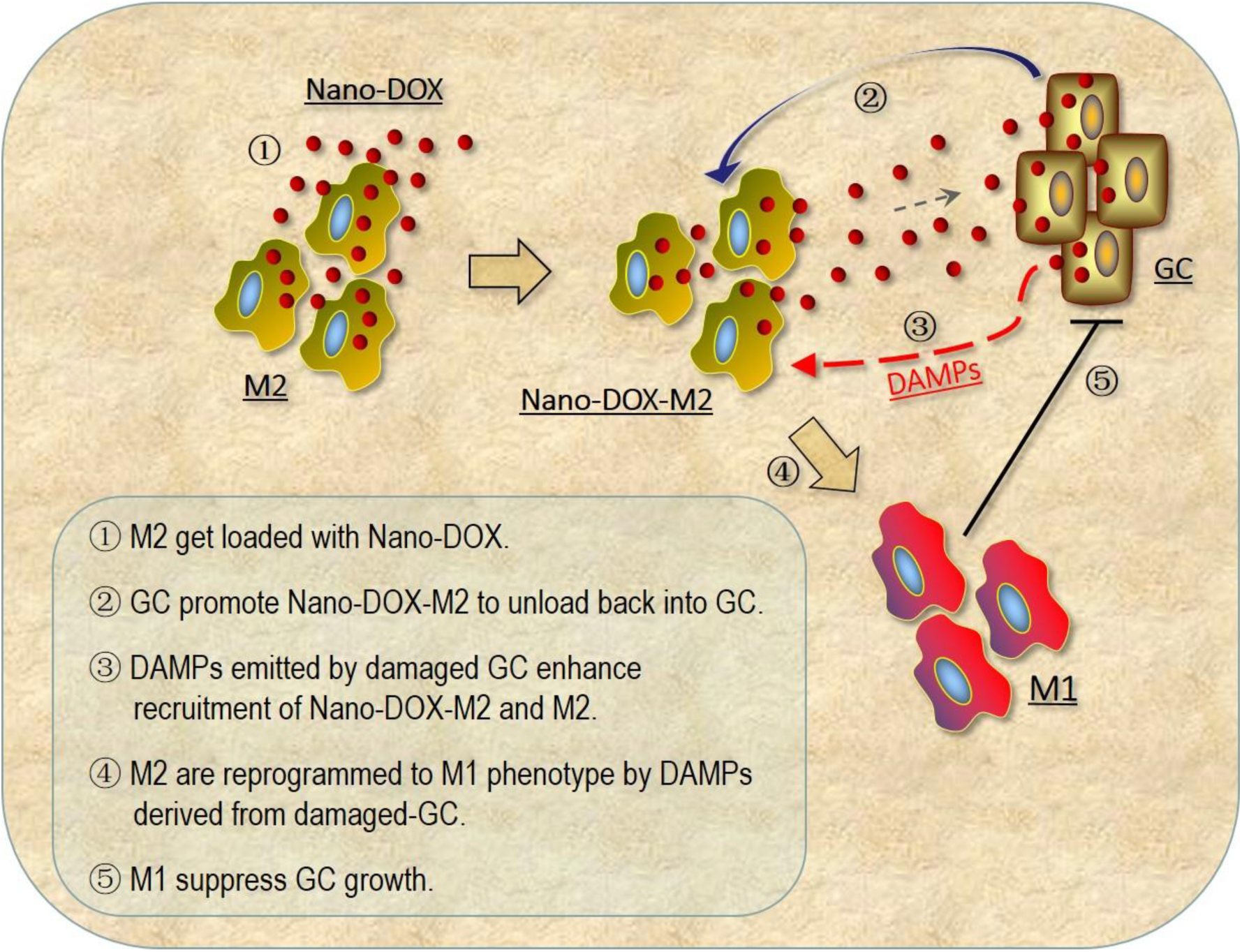
(Also as TOC Graphic) GC have close cross-talks with their TAM, which could be exploited on two dimensions for therapeutic modulation of the GBM microenvironment. First, TAM serve as a carrier and reservoir to release drugs to the cancer cells and unloading of drug-loaded TAM is promoted by the cancer cells. Second, damaged GC recruit more TAM, non-loaded as well as drug-loaded, and reprogram them toward an anti-tumor phenotype, wherein GC-derived danger signal i.e. the DAMPs may play a central role.

Second, damaged GC recruit more TAM, non-loaded as well as drug-loaded, and reprogram them toward an anti-tumor phenotype, wherein GC-derived danger signals i.e. the DAMPs may play a central role. On one hand, the enhanced recruitment of drug-loaded TAM by damaged GC may create a positive feedback loop that favors therapy as greater GC damage would result in more TAM-mediated drug delivered in the tumor. On the other hand, due to TAM’s central role in orchestrating the tumor immune microenvironment, reprogramming of the TAM has been increasingly appreciated for its therapeutic value in modulating tumor microenvironment and thereby in cancer treatment. Tumor cell-derived DAMPs are key effector molecules involved in Mø activation and implicated in the therapeutic efficacy of certain anti-cancer drugs like mitoxantrone, oxaliplatin, and anthracyclines to which DOX belongs [33, 36, 42]. Our work compellingly demonstrates Nano-DOX to be a powerful tool for TAM-reprogramming owing to its DAMPs-inducing property which is more potent than DOX (Fig. 4 – Fig. 6, Fig. 13). DAMPs are a variety of signal molecules with different intracellular sources and release mechanisms. For example, CRT mainly comes from the endoplasmic reticulum and cytoplasm; HSP90 is a cytosolic molecular chaperone; and HMGB1 is a chromatin protein in the nucleus under normal condition. The mechanisms by which Nano-DOX induce DAMPs emission are an intriguing question. DOX as an established DAMPs release inducer may provide a point of reference, but Nano-DOX does not necessarily conform to DOX’s pattern. For instance, Nano-DOX increased HMGB1 release from GC whereas DOX suppressed HMGB1 release (Fig. 5). As mentioned earlier, our unpublished data showed that while DOX essentially induce apoptosis, Nano-DOX is a potent inducer of autophagy and there is mounting evidence that autophagy regulates release of DAMPs such as ATP, HMGB1, IL-1β and etc [43]. Further work is ongoing to investigate the possible link between Nano-DOX induced autophagy and DAMPs emission. Interestingly but not surprisingly, Nano-DOX also induces DAMPs from Mø (Fig. 4A, Fig. S4) which suggests Nano-DOX may have wider implications in the modulation of tumor immune microenvironment as Mø-derived DAMPs may potentiate Mø’s immune function. For example, Mø are reported to use their own CRT to guide recognition and phagocytosis of adjacent tumor cells [44].

The primary novelty of this work is the proposition to modulate the GBM immune microenvironment through harnessing the cross-talk between GC and their associated macrophages with a nano-drug i.e. Nano-DOX. The anti-GBM efficacy of this strategy should, ideally, be demonstrated in vivo on GBM models. However, it would not be possible to tell whether the therapeutic effect comes from Nano-DOX’s cytotoxicity or from its modulation of the GBM microenvironment. Comparison of Nano-DOX-M2 against Nano-DOX of the same dosage may help to solve the problem. As shown in Fig. 11 & Fig. 12, however, Nano-DOX per se could hardly penetrate into the orthotopic tumor tissue, thus direct comparison of the anti-GBM effects of Nano-DOX-M2 and Nano-DOX in vivo is not practical. Mø-mediated delivery of Nano-DOX in the GBM is another key notion of this work. We propose herein that Mø not only serve as drug carriers but can also be modulated as effector cells albeit Mø-mediated drug delivery in tumor or the central nervous system is not an original idea. In consideration of the above, instead of demonstrating Nano-DOX’s anti-GBM effect in vivo, we have chosen to ① compare the DAMPs-emission-inducing capacity of Nano-DOX-M2 against Nano-DOX and DOX (Fig. 13) and ② prove the M2-M1 phenotype shift of Mø induced by Nano-DOX in the orthotopic GBM xenografts (Fig. 14). Results obtained and presented in this manuscript are adequate and constitute a logical chain of evidence substantiating our propositions. A separate study is underway assessing the in vivo anti-GBM efficacy of Nano-DOX delivered by monocytes.

It should be noted that organs such as the heart and liver appear able to accumulate Nano-DOX and Nano-DOX-mBMDM2, which may give rise to cardiotoxicity and liver toxicity. This concern which might affect the translational potential of Nano-DOX, along with their in vivo pharmacokinetics, is being addressed in our further work.

## 5. Conclusion

Nano-DOX provides a novel approach to modulating the glioblastoma immune microenvironment through harnessing the cross-talk between cancer cells and their associated macrophages.

## Acknowledgments

This research was supported by the National Natural Science Foundation of China (No. 81671818), Natural Science Foundation of Hubei Province, China (No.2015CFB403) and Science and Technology Program of Wuhan, China (No. 2017060201010148).

## Supporting Information

Additional data are supplied as Supporting Information.

**Fig. S1:**
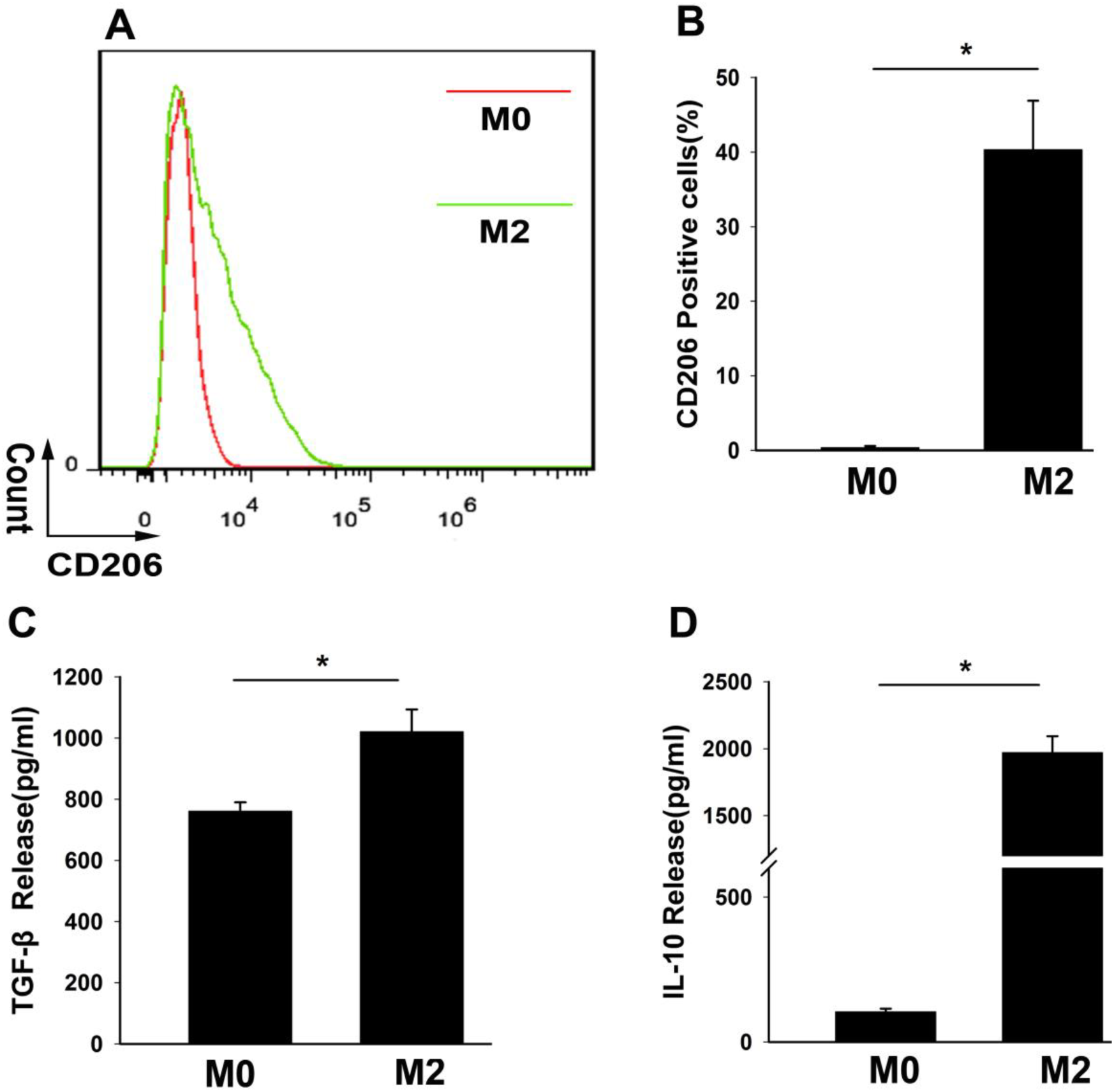
Characterization of M2 by cell surface marker and cytokine release. A: CD206, cell surface marker for type-2 Mø activation, was analyzed by immunofluorescent staining and FACS. B: Geometric means were used to quantify fluorescence intensity. C and D: Cytokine levels of TGF-β and IL-10 were assayed by ELISA. Values were means ± SD (n=3, * p < 0.05). Unpolarized Mø (M0) were used as control.

**Fig. S2:**
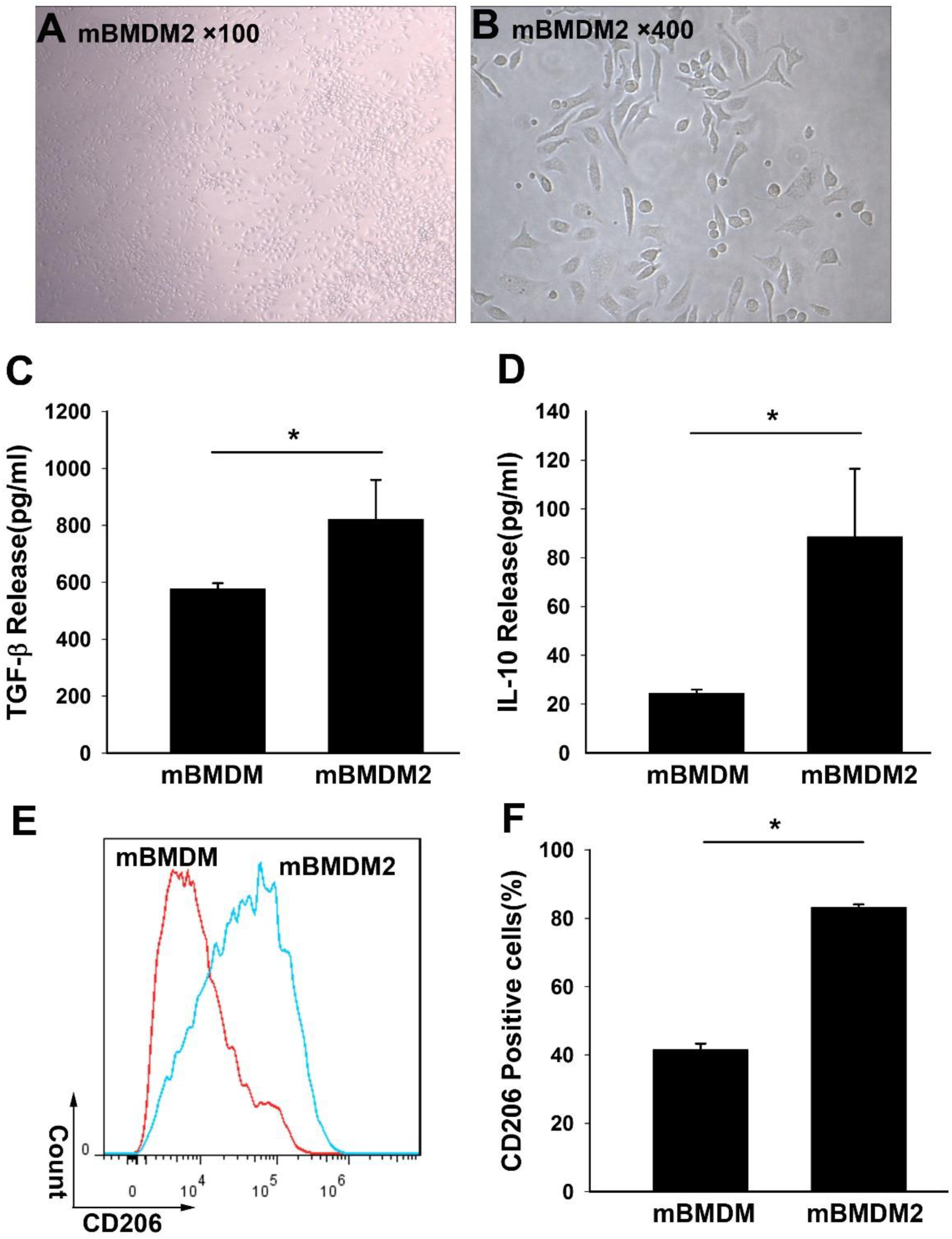
Characterization of mBMDM2 by cell surface marker and cytokine release. A and B: Visuals of mBMDM2 were captured on day 7 of polarization. C and D: Cytokine levels of TGF-β and IL-10 were assayed by ELISA. E: Cell surface marker CD206 was analyzed by immunofluorescent staining and FACS. F: Geometric means were used to quantify fluorescence intensity. Values were means ± SD (n=3, * p < 0.05). Unpolarized Mø (M0) were used as control.

**Fig. S4:**
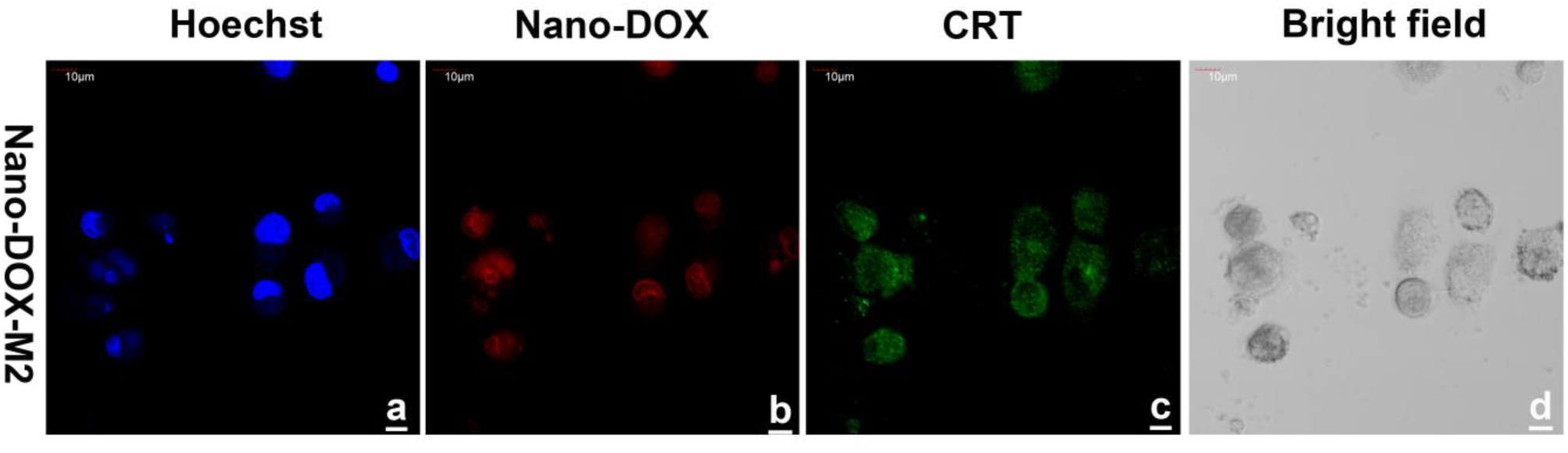
Single-cultured M2 were treated with 2.5 μg/mL of Nano-DOX for 24h. Cell surface exposure of CRT was detected by immunofluorescent staining. Green fluorescence is CRT staining; red fluorescence comes from Nano-DOX and blue fluorescence is nuclear counterstaining.

**Fig. S5:**
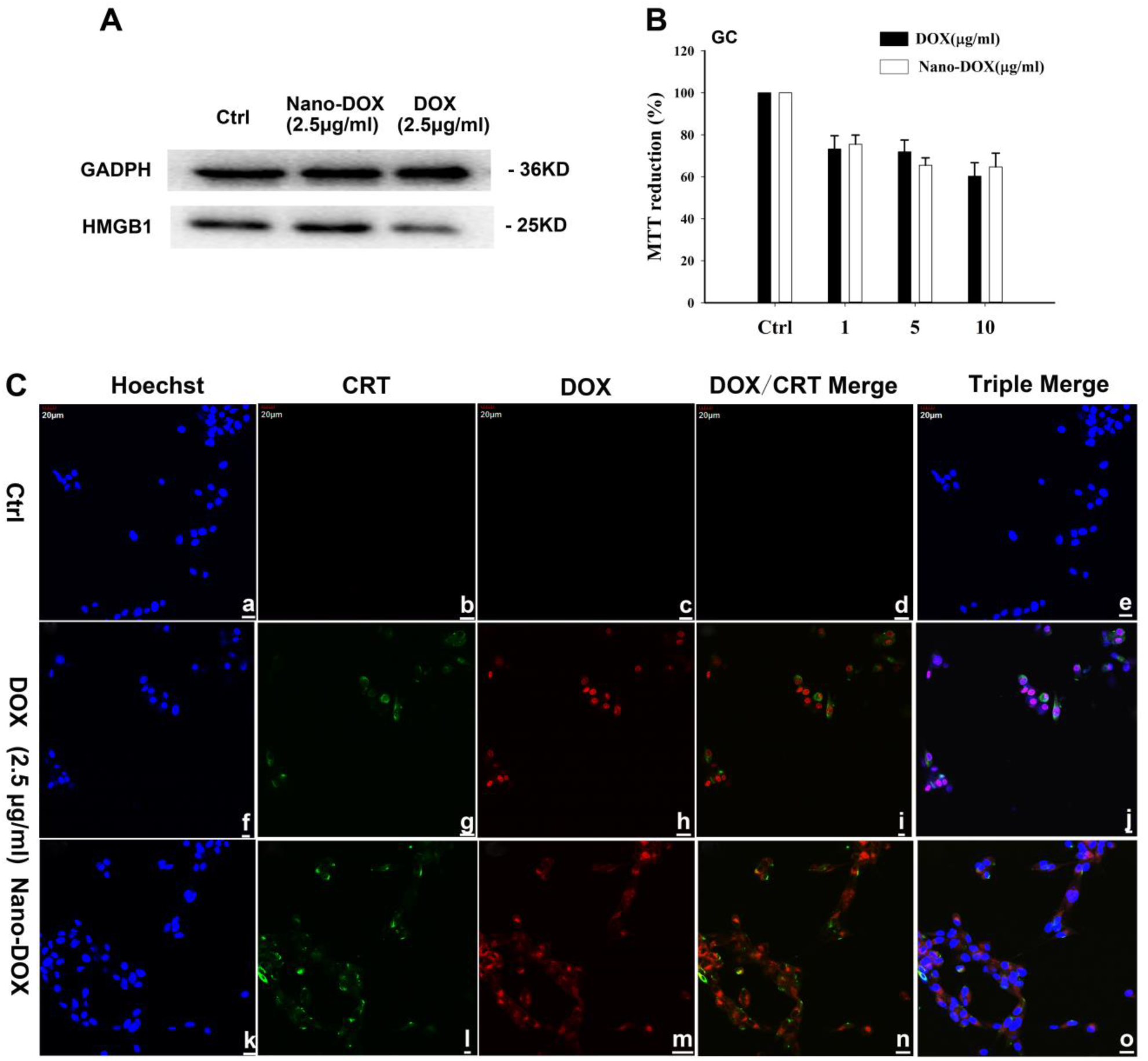
A: GC were treated with DOX or Nano-DOX (2.5μg/mL) for 24h and expression of HMGB1 were detected by Western analysis. B: GC were treated with DOX or Nano-DOX and cell viability was assayed by MTT reduction. C: Triple merge of fluorescence of Hoechst, CRT staining and DOX/Nano-DOX were supplemented in Fig. 5A.

**Fig. S7:**
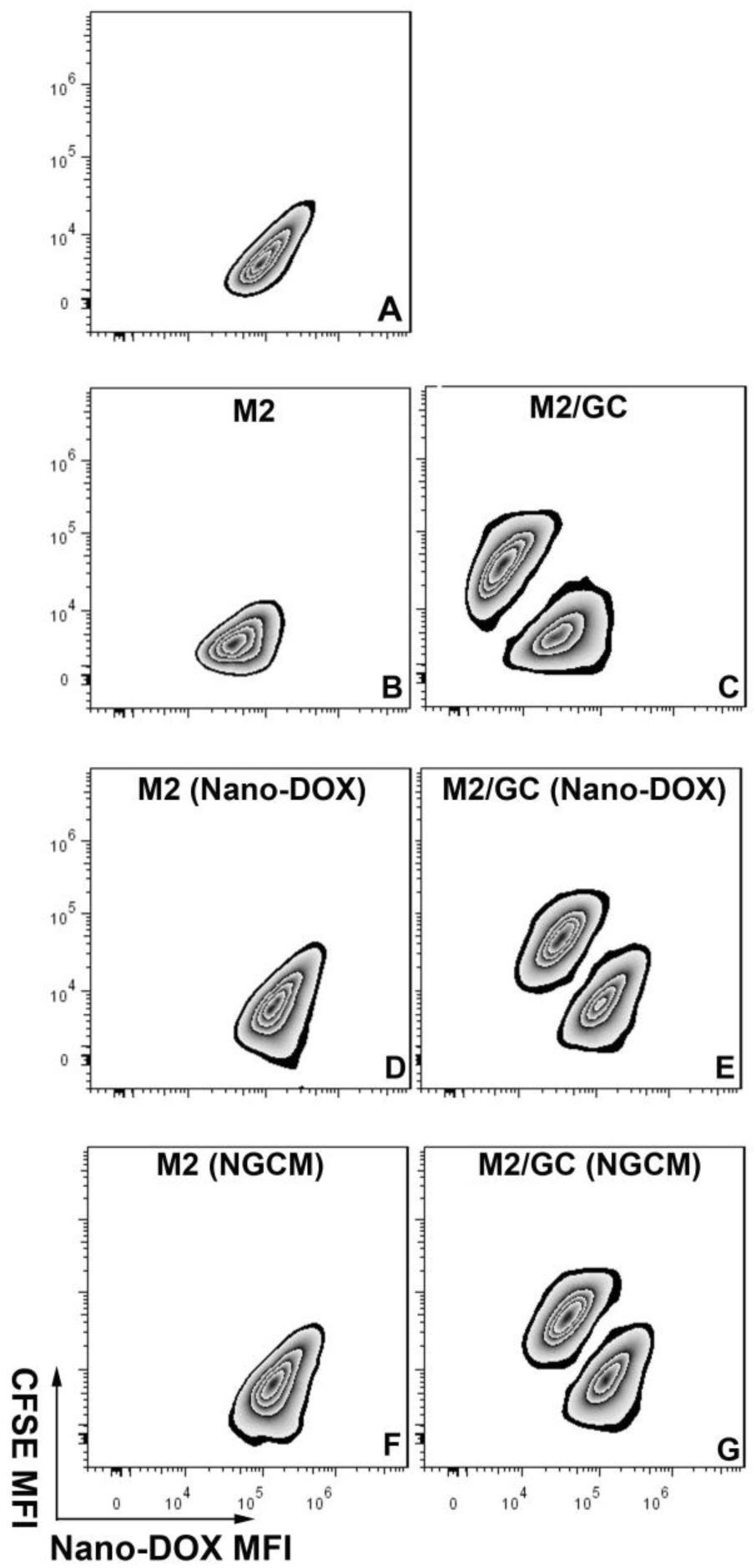
Representative FACS zebra dot plots for Fig. 6 H. Nano-DOX-M2 (A) were mixed-cultured with CFSE-labelled GC for 24 h in RM (C), RM with Nano-DOX (E) or NGCM (G). Single-cultured Nano-DOX-M2 (B, D & F) served as controls.

**Fig. S8:**
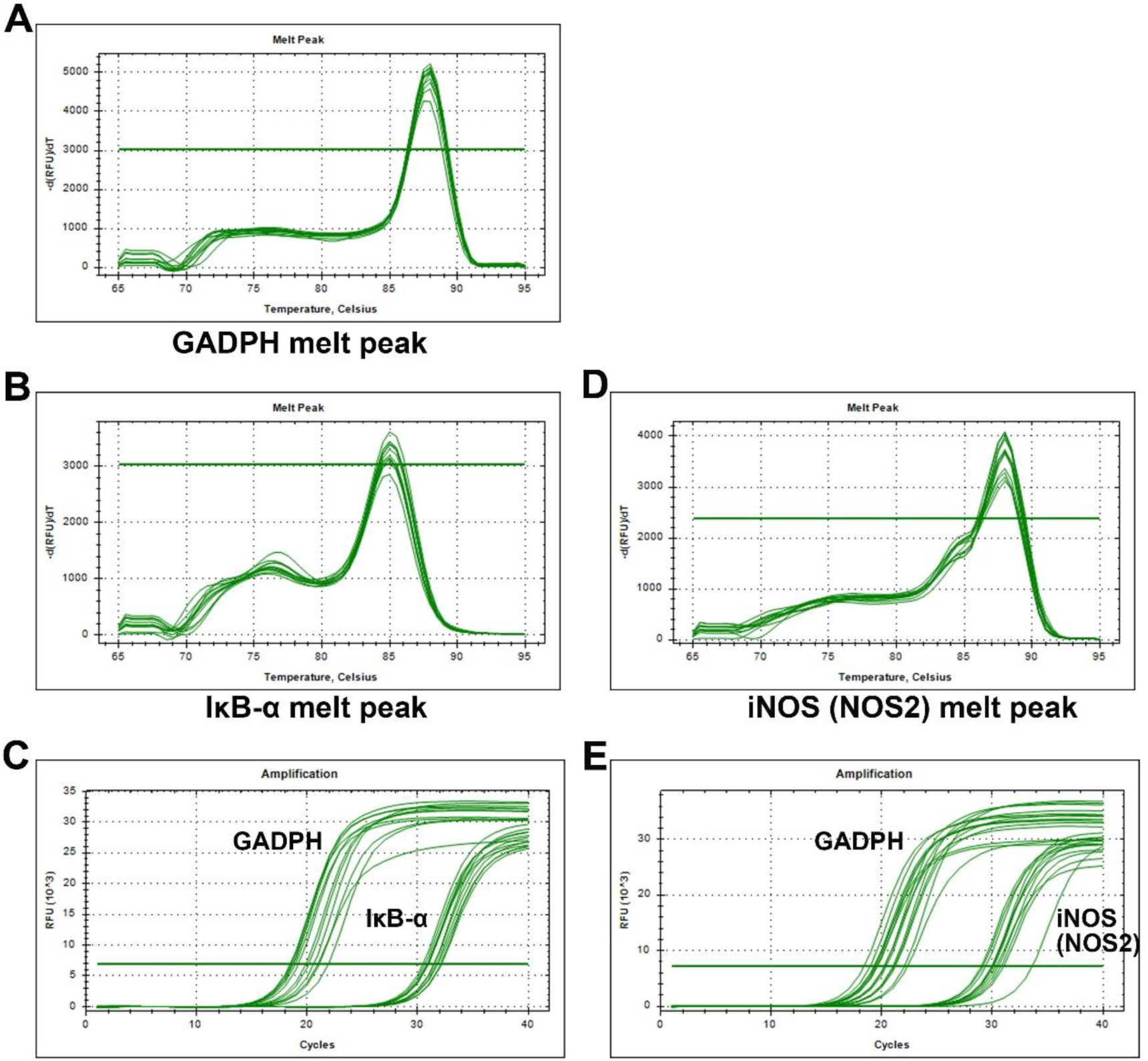
Melting curves and amplification curves for Fig. 8 D and E. A, B and D: Melting curves of GADPH, IκB and iNOS. C and E: amplification curves of GADPH, IκB and iNOS.

**Fig. S9:**
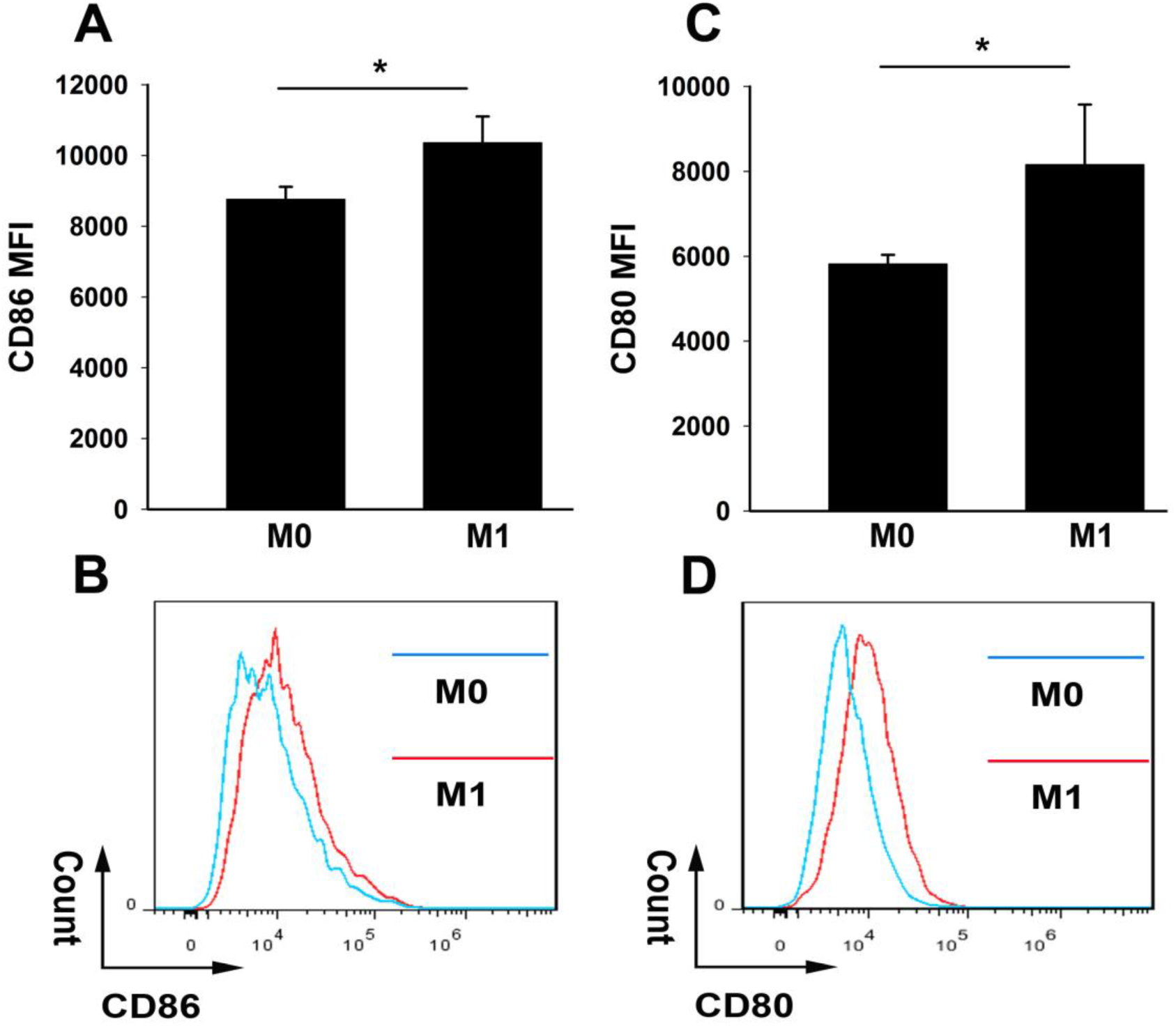
CD80 and CD86 expression in M1 as the positive control for Fig. 8 C & E. Cell surface markers CD86 (A, B) and CD80 (C, D) expression in M1 were analyzed by immunofluorescent staining and FACS. Geometric means were used to quantify fluorescence intensity. Unpolarized Mø (M0) were used as controls.

**Fig. S9-2:**
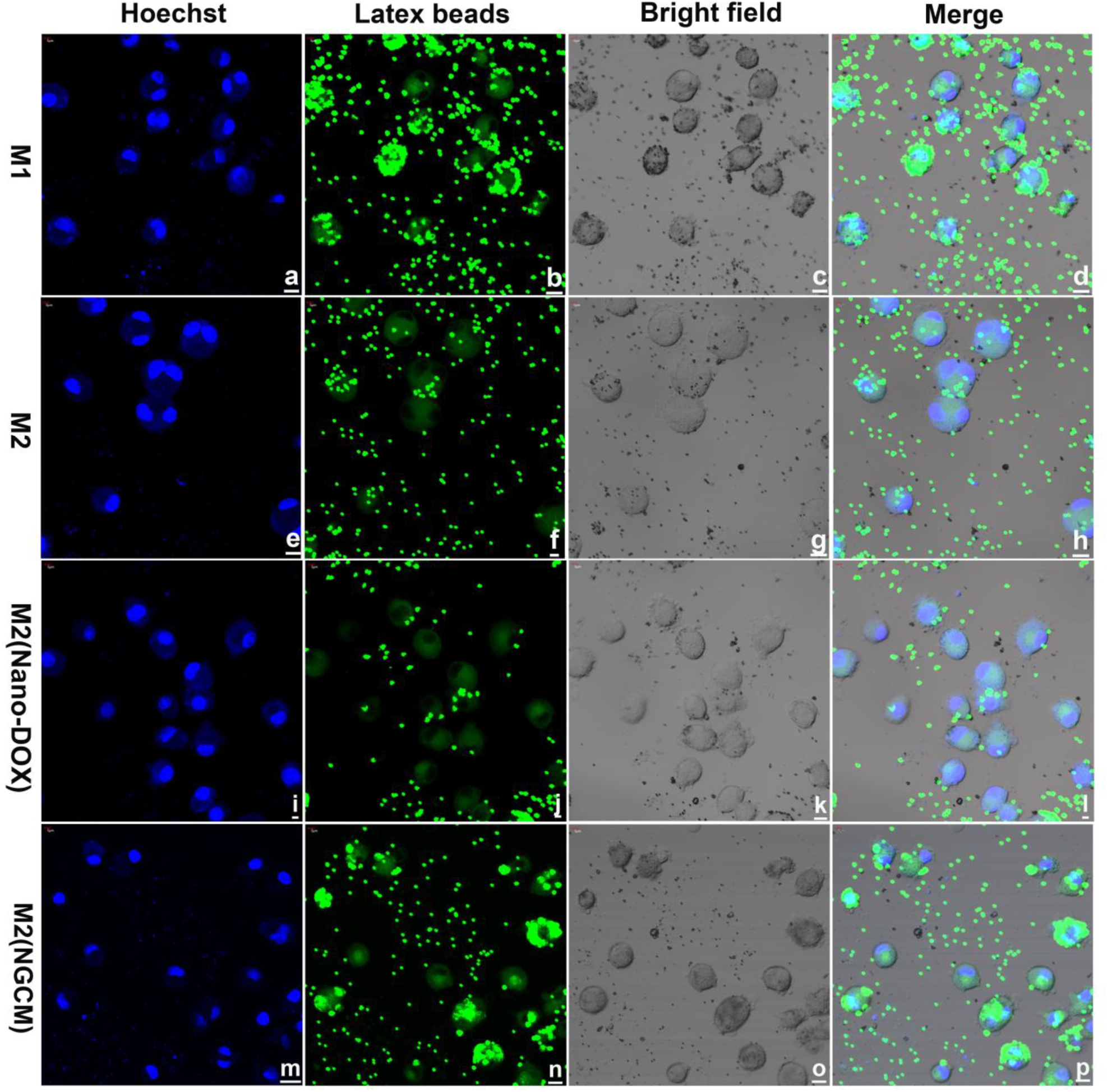
A comprehensive version of Fig. 8 H with more controls. Green fluorescence is FITC-latex beads counterstaining.

**Fig. S10:**
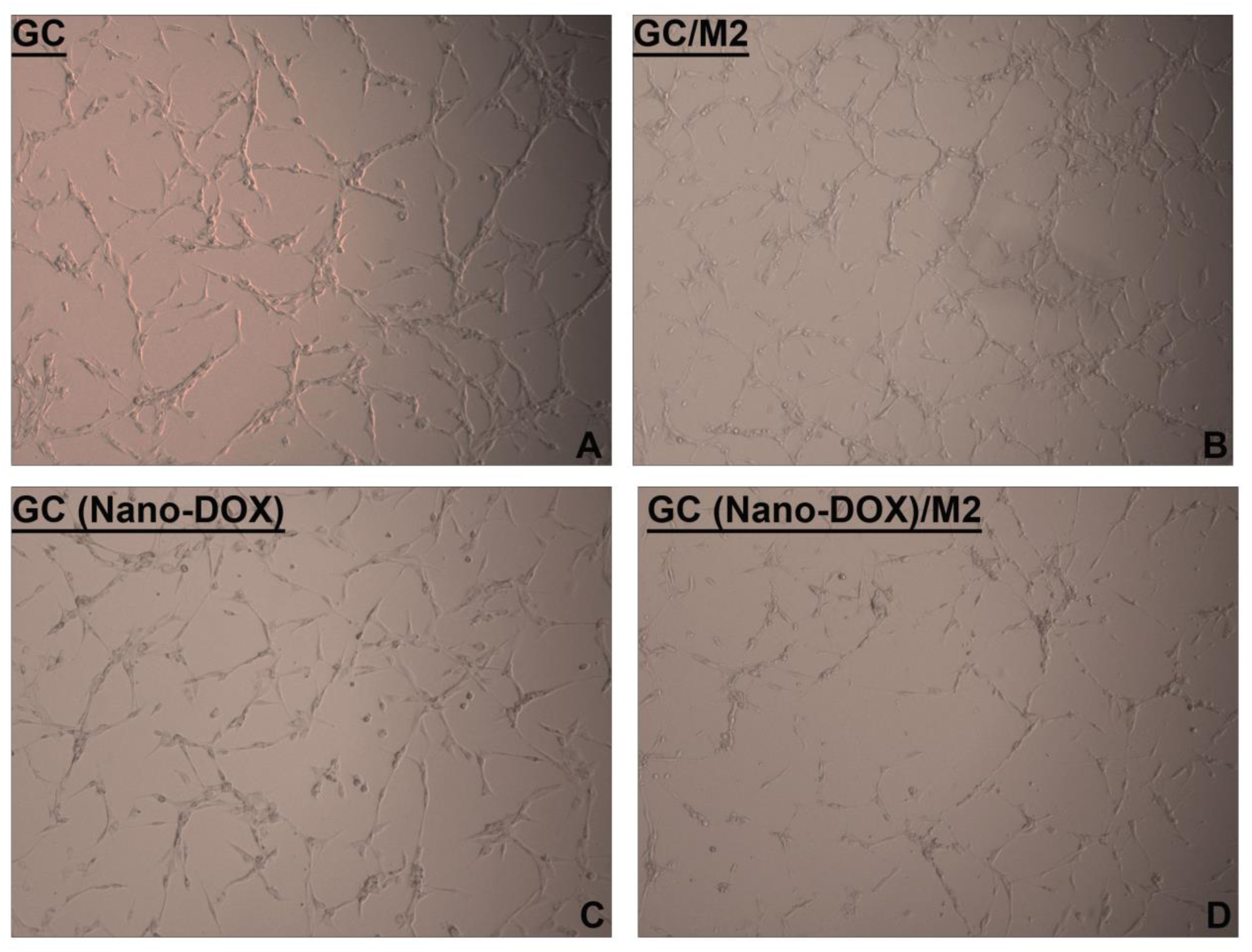
Visuals of GC growth upon CCK-8 test in Fig. 9 B. A: GC were single cultured for 24h. B: GC were co-cultured with M2 in a Transwell for 24h. C: GC were singled cultured in the presence of 2.5 μg/mL of Nano-DOX for 24h. D: GC were co-culture with M2 in the presence of 2.5 μg/mL of Nano-DOX in a Transwell for 24 h.

**Fig. S11:**
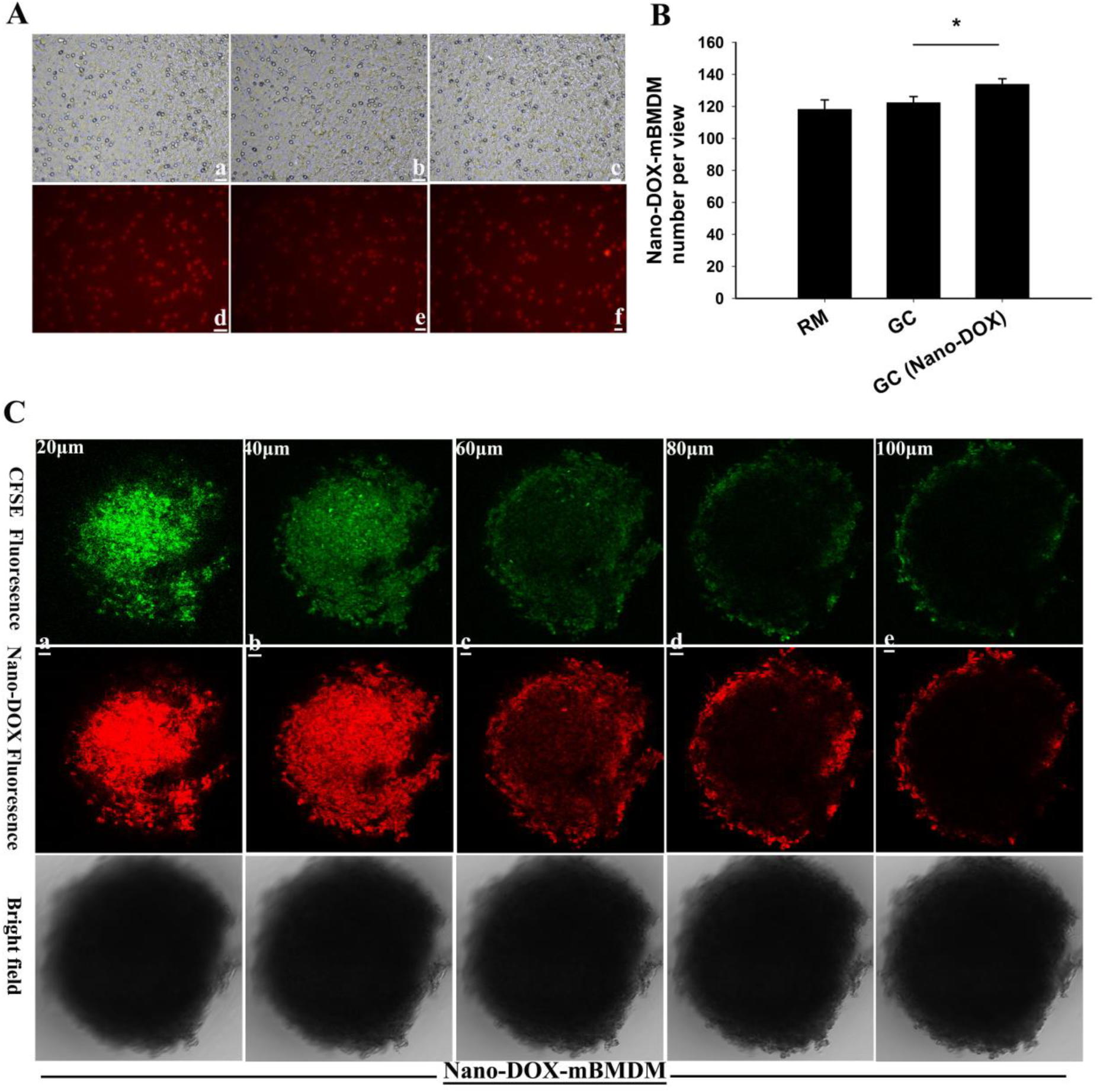
Demonstration of functional viability of Nano-DOX-mBMDM2 by Transwell chemotaxis assay and 3-D GC spheroid infiltration assay. A: Nano-DOX-mBMDM2 that were attracted by GC to the underside of the insert bottom membrane were photographed. B: Cells in 5~8 fields of view were counted. C: Nano-DOX-mBMDM2 were mixed-cultured with GC spheroids for 12 h and the spheroids were photographed at different depths using a laser confocal microscope. Nano-DOX-mBMDM2 were prepared by incubating mBMDM2 with 2.5 μg/ml of Nano-DOX for 24 h.

**Fig. S12:**
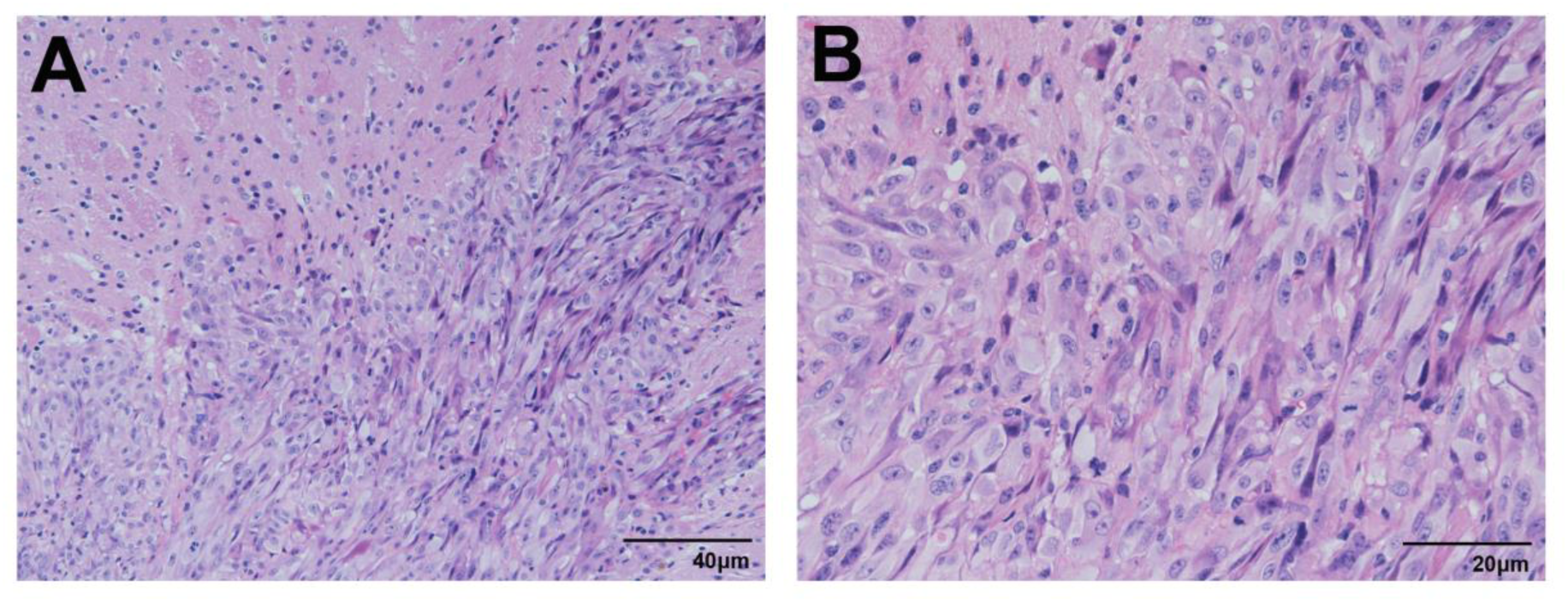
Representative HE staining of orthotopic GC xenograft tissues from mice. A: 200×magnification. B: 400×magnification.

